# Task-specific topographic organization of neural activity in the primate lateral prefrontal cortex

**DOI:** 10.1101/2024.05.10.591729

**Authors:** Jinkang Derrick Xiang, Megan Roussy, Benjamin Corrigan, Roberto A. Gulli, Rogelio Luna, Maryam Hasanzadeh Mofrad, Lyle Muller, Jörn Diedrichsen, Taylor W. Schmitz, Julio Martinez-Trujillo, Marieke Mur

## Abstract

Neurons in the primate lateral prefrontal cortex (LPFC) flexibly adapt their activity to support diverse cognitive demands. Whether this flexibility is accompanied by changes in the spatial organization of neural population activity remains unclear. Here, we characterized the functional topography of LPFC neural activity in awake, behaving macaques performing three distinct cognitive tasks. Using chronically implanted multi-electrode arrays and topographic similarity analysis (TSA), we found that LPFC topographies were reproducible within tasks but less consistent across tasks. We then separated neural activity into task-tuned and trial-to-trial residual components to determine the basis of these effects. Task-tuned topographies mirrored the overall pattern: they were less consistent across than within tasks, whereas residual topographies remained comparatively stable across tasks. Moreover, task-tuned and residual topographies showed opposing patterns of consistencies. Within tasks, task-tuned topographies were more consistent than residual topographies; across tasks, the pattern reversed. This dissociation suggests a reproducible spatial organization of task-related activity that reconfigures across tasks. Nevertheless, task-tuned topographies remained significantly correlated across tasks, revealing a shared component of spatial organization. Across all three tasks, topographies exhibited a common spatial scale comparable to that of patterned afferent inputs to LPFC reported in previous anatomical tracing studies. Together, these findings suggest that LPFC functional organization combines a shared spatial scaffold with selective, task-dependent reconfiguration of task-tuned population activity.

## 1. Introduction

Flexibility is a defining property of the higher-order cognitive functions supported by the primate lateral prefrontal cortex (LPFC). Unlike neural populations in primary sensory areas, where activity is strongly shaped by stimulus features such as frequency and orientation (Hubel and Wiesel, 1968, Merzenich and Brugge, 1973), LPFC neurons adapt their activity according to rules, behavioral context, and feedback, even when sensory inputs are held constant (Duncan, 2001, Freedman et al., 2001, Miller and Cohen, 2001, Lennert et al., 2011, Lennert and Martinez-Trujillo, 2013). This flexibility is shaped in part by the diverse inputs received by LPFC neurons, which are more heterogeneous than those of sensory neurons (Goldman-Rakic, 1988, Fuster, 2015). Consistent with this connectivity, LPFC neurons often exhibit selectivity for combinations of task features, such as firing maximally for a specific conjunction of rule, context, and feedback (Rigotti et al., 2013, Fusi et al., 2016).

The heterogeneous and context-dependent selectivity of individual LPFC neurons creates challenges for understanding their functional organization. In visual cortex, neurons with similar stimulus preferences are often spatially clustered, giving rise to functional organizations that can be delineated using stimulus mapping techniques (Wandell et al., 2007). Mapping visual field location (Brewer et al., 2002), stimulus orientation (Fang et al., 2022), or object category (Bell et al., 2009, Bao et al., 2020) reveals spatially organized populations with distinct feature preferences in striate and extrastriate cortex. These stimulus-tuned populations can remain spatially organized over days, weeks, and months (Margolis et al., 2012, McMahon et al., 2014, Cossell et al., 2015), forming stable topographic maps. Such maps are organized at a columnar spatial scale (Hubel and Wiesel, 1968, Tanaka, 1996) that can be detected using multi-electrode arrays and functional magnetic resonance imaging (fMRI) (Brewer et al., 2002, Yacoub et al., 2008, Sato et al., 2013). Whether analogous topographies exist for task-related activity in the LPFC remains unclear (Constantinidis et al., 2001, Markowitz et al., 2015, Masse et al., 2017, Bullock et al., 2017, Leavitt et al., 2018). In particular, it is unknown whether LPFC topographies are stable over time (Driscoll et al., 2017, Muysers et al., 2024), reorganize according to task demands, and operate at a spatial scale comparable to topographic organization in other cortical areas. Testing these possibilities has been challenging because the task space sampled in individual experiments is typically limited in both complexity and diversity (Yang et al., 2019a). Primate electrophysiology studies often examine a single task or task features that are insufficiently distinct, and few have examined changes in the spatial organization of population activity across recording sessions or tasks.

If task-dependent functional topographies exist in LPFC, two properties are particularly important for understanding their contribution to flexible cognition. First, a functional topography should be sufficiently stable within a task to provide a reproducible spatial organization of task-related activity. Second, it should be sufficiently adaptable across tasks to allow a common LPFC network to support different cognitive demands. Thus, a central question is whether LPFC topography primarily reflects a stable spatial architecture or is reorganized according to task demands. These possibilities are not mutually exclusive. A stable spatial architecture could provide a scaffold for reliable task-specific population codes while allowing their spatial organization to be selectively reconfigured across tasks.

Here, we tested these possibilities using a unique dataset of multi-electrode array recordings from the LPFC of two macaques trained to perform three distinct cognitive tasks (Luna et al., 2019, Roussy et al., 2021, Corrigan et al., 2022). The tasks sampled a diverse task space by differentially engaging cognitive demands, embedding stimuli within rich behavioral contexts, and using virtual-reality environments to promote engagement of the LPFC (Doucet et al., 2016). The same tasks were performed across recording sessions spanning multiple days, allowing us to assess the stability of population topography over time and its reorganization across tasks.

To identify task-specific functional topographies, we adapted representational similarity analysis (RSA) (Kriegeskorte et al., 2008a) to develop topographic similarity analysis (TSA). RSA is widely used in fMRI (Kriegeskorte et al., 2008b, Mur et al., 2013) and electrophysiology studies (Kiani et al., 2007, Freiwald and Tsao, 2010) to characterize the distributed response patterns across measurement channels, such as voxels or electrodes, evoked by individual stimuli. Comparing these response patterns across stimuli yields an estimate of representational geometry that is independent of the specific measurement channels used to obtain the responses (Kriegeskorte and Kievit, 2013) (Fig. 1). TSA reverses this perspective by focusing on the transpose of the response matrix: the response profile of each measurement channel across multiple task features (Kiani et al., 2015). Comparing these profiles across channels yields an estimate of their functional topography while abstracting from the specific task features used to characterize the response profiles (Fig. 1).

**Figure 1:**
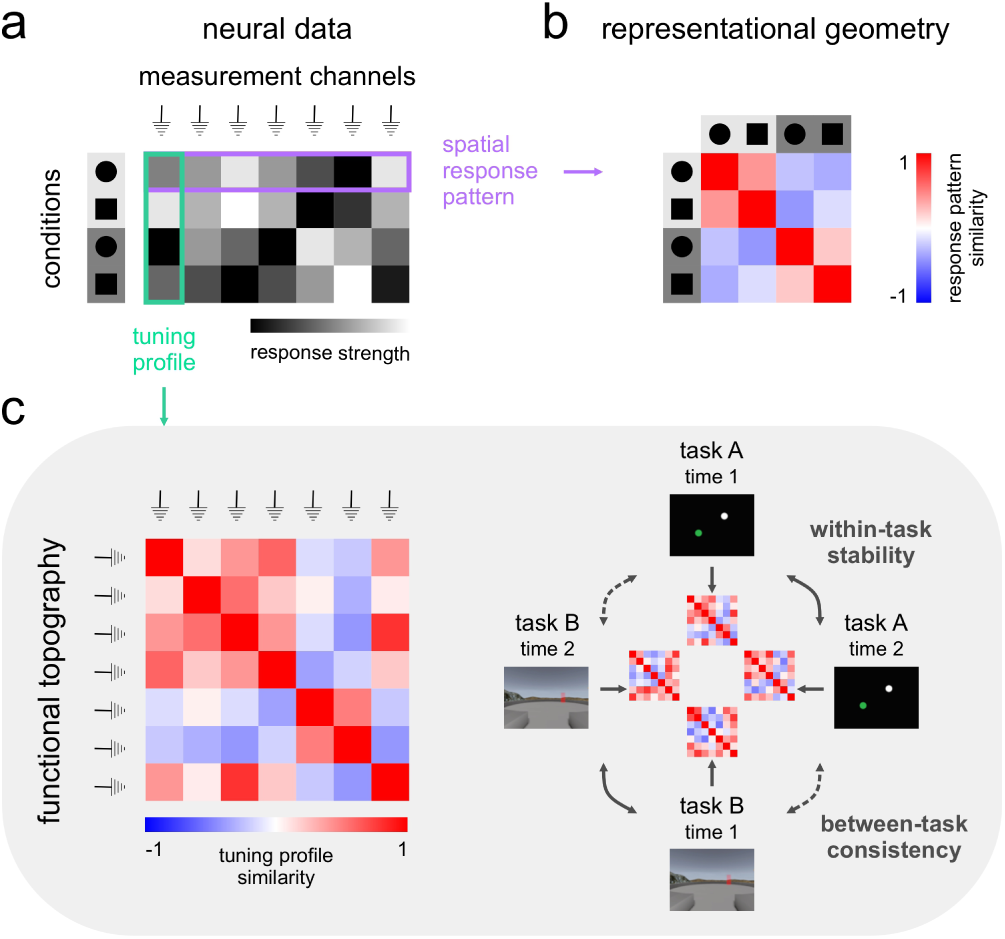
Topographic similarity analysis. To test for task-specific functional topographies in the LPFC, we adapt representational similarity analysis (RSA) to topographic similarity analysis (TSA). a) Simulated neural data across experimental conditions (rows) and measurement channels (columns). Higher spike rates in a given channel are denoted by lighter shades. **b**) RSA abstracts from measurement channels to infer representational geometry. **c**) TSA abstracts from experimental conditions to infer functional topography. TSA allows for quantitative comparison of functional topographies across time and tasks.

TSA offers two advantages for characterizing functional organization in the LPFC. First, it incorporates the full response profile of each channel across multiple task features, allowing spatial organization to reflect context-dependent combinations of neural selectivity rather than tuning to a single predefined feature. Second, by comparing channel-level response profiles through their pairwise similarities, TSA provides a framework for characterizing functional topography across tasks, even when the tasks involve different features (Xiang et al., 2025).

We found that LPFC topographies of task-tuned activity were stable within tasks but selectively reorganized across tasks. Task-tuned topographies were more stable across recording sessions than topographies of trial-to-trial residual activity, but showed greater task-dependent change. Despite this reorganization, task-tuned topographies retained a significant shared component across tasks. Finally, the task-specific topographies exhibited a common spatial scale that was consistent with the scale of patterned afferent projections to LPFC described in previous anatomical tracing studies (Goldman-Rakic, 1984, Goldman-Rakic and Schwartz, 1982, Leichnetz et al., 1981). These findings suggest that LPFC functional organization comprises a spatial scaffold that is sufficiently stable to support reproducible task-related activity while remaining adaptable to changing cognitive demands (Xiang et al., 2026, Nettekoven et al., 2026).

## 2. Results

### 2.1. Estimating the topography of LPFC neural activity across time and tasks

The same two rhesus macaque monkeys (monkey B and monkey T) each performed three cognitive tasks (Fig. 2a-c). The first task was an oculomotor delayed response task (ODR) (Luna et al., 2019). The second task was a visuospatial working memory task (VWM) deployed in a virtual-reality environment with naturalistic scenes for stronger attentional engagement (Roussy et al., 2021, Doucet et al., 2016). The third task was a context-dependent decision making task (CDM), which was also deployed in a virtual-reality environment (Corrigan et al., 2022). Altogether, these three tasks engage a wide spectrum of cognitive functions, including working memory, visuospatial attention, context-dependent decision making and motor planning.

**Figure 2:**
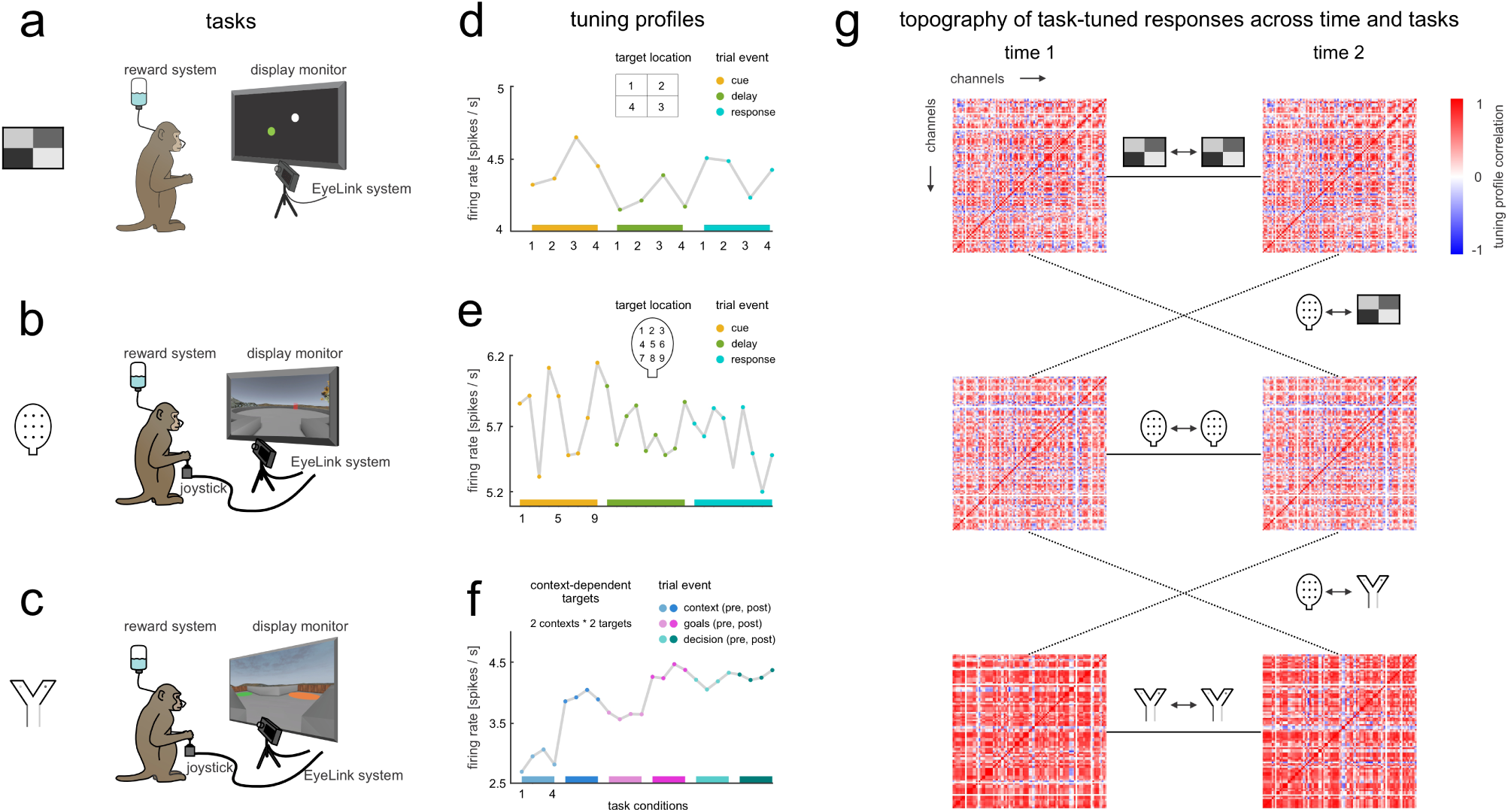
Comparing task-tuned functional topographies in the LPFC across time and tasks. **a)** Oculomotor delayed response task (ODR). Monkeys fixated a point on the screen. A visual cue appeared, then disappeared. After a delay, the monkeys saccaded toward the remembered target location. **b**) Visuospatial working memory task (VWM). A visual cue appeared in one of nine target locations in a virtual arena, then disappeared. After a delay, the monkeys navigated toward the remembered target location using a joystick. **c**) Context-dependent decision making task (CDM). Monkeys navigated through an X maze using a joystick. The texture of the corridor walls indicated the decision context for target selection. **d-f**) Example tuning profile for a single channel in the ODR (d), VWM (e), and CDM (f) tasks. Legends indicate the task features that composed the unique task space. **g**) Schematics for TSA of task-tuned responses across time (solid lines) and tasks (dashed lines). Matrix elements are Pearson correlations of tuning profiles between channel pairs.

We recorded the responses of neurons in layers II/III of LPFC areas 8A and 9/46, dorsal and ventral to the principal sulcus, using 96-channel multi-electrode Utah arrays (see Supplementary Fig. 1 for array placement). Each array covered a 4 mm *×* 4 mm cortical area with 10 *×* 10 electrodes (*∼*0.4 mm spacing). Both monkeys performed multiple measurement sessions for each task on separate days (Supplementary Fig. 2). In each session, we simultaneously recorded neural activity from multiple channels on each array (see Supplementary Table 1 for details on active channels). Action potential times were extracted and synchronized to task events. To enable spatial mapping of response preferences across the array, we pooled spiking activity from putative units identified on the same recording channel, summing their activities to create multi-unit activity. Units measured by the same channel exhibited similar response preferences (Supplementary Fig. 3). The pooled activity reflects the activity of subpopulations of neurons within the area covered by each array channel.

To characterize the spatial organization of population codes in the LPFC, we first computed task tuning profiles, one for each channel in each session in each task (Fig. 2d-f). Task tuning profiles are vectors that store the firing rates of a channel to the experimental conditions. The time windows for estimating spike rates vary from a few hundred milliseconds up to a few seconds (see Methods for details). These time windows were determined based on task structure, monkey behavior and population decoding results. Next, we computed a channel-by-channel tuning similarity matrix for each array in each session in each task (Fig. 2g). Each element of the matrix represents the Pearson correlation of tuning profiles between a channel pair. The matrix as a whole reflects the similarity of task tuning for all channel pairs, thus capturing the functional topography of task-tuned LPFC activity. This characterization enables quantitative comparison of functional topographies across time and tasks (Figs. 1, 2g). For comparative purposes, we also analyzed trial-to-trial fluctuations around the trial averages that define the tuning profiles (see Methods). Topographies based on these spontaneous fluctuations, or residuals, are expected to be consistent across tasks (Cole et al., 2014, Kiani et al., 2015).

### 2.2. The topography of LPFC neural activity is stable over time but adaptive across tasks

We first asked whether the functional topography of LPFC neural activity is stable over time and consistent across tasks (ODR, VWM, and CDM). We used TSA to quantify topographic consistency as the correlation between channel-by-channel similarity matrices across sessions (Fig. 2g). Correlations between sessions within the same task quantify the stability of topography over time, whereas correlations between sessions from different tasks quantify its consistency across tasks. For each array in each monkey, topographies were estimated from the overall time series of each channel, independent of task condition. This definition allowed us to assess topographic organization without introducing differences arising from the specific task conditions sampled across sessions.

To minimize the effects of changes in array position across recording sessions, we restricted both within-and between-task comparisons to session pairs separated by no more than 20 days. Because some tasks were recorded more than 20 days apart (Supplementary Fig. 2), each monkey contributed between-task comparisons for two of the three tasks. We focus below on the ventral array in both monkeys because recordings from the dorsal array in mT had a lower signal-to-noise ratio (see Methods).

#### Topographies were stable over time within a task

For within-task comparisons, topographic consistency was significantly greater than zero for all task pairs examined (one-sided permutation tests, *p <* 0.05; Fig. 3a,b). The mean consistency was *r* = 0.50 for VWM-VWM and *r* = 0.21 for CDM-CDM in mB, and *r* = 0.60 for VWM-VWM and *r* = 0.14 for ODR-ODR in mT. Thus, despite changes in recording session, neural populations with similar response structure tended to retain similar relative organization within a given task. The dorsal array in mB showed a similar pattern (Supplementary Fig. 4).

**Figure 3:**
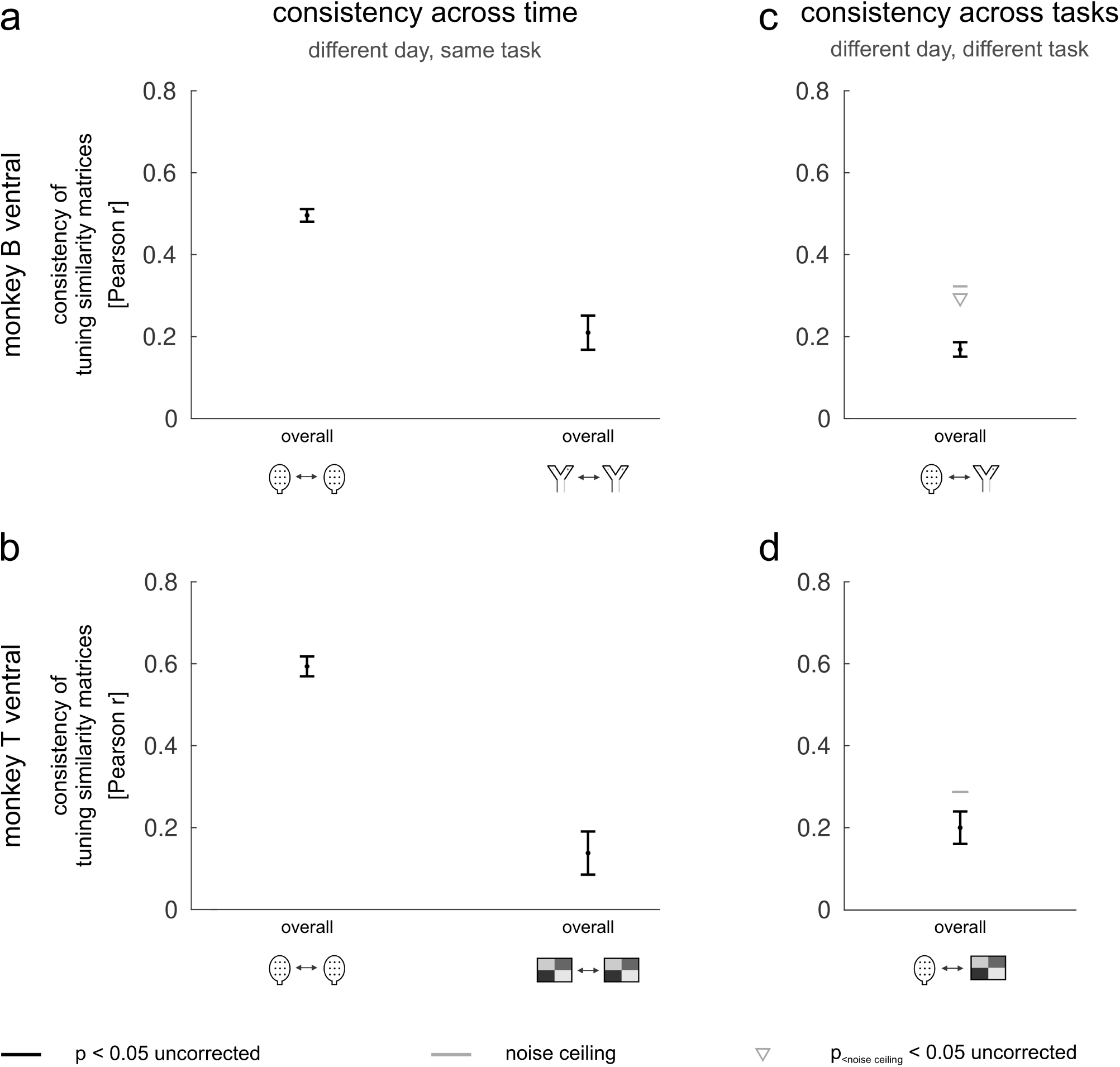
The topography of LPFC neural activity is stable over time but adaptive across tasks. **a-b)** Consistency of functional topographies across sessions within the same task for mB (a) and mT (b). **c-d**) Consistency of functional topographies across sessions between different tasks for mB (c) and mT (d). Functional topographies were estimated from the overall time series of each channel, independent of task condition. Consistency was quantified as the correlation between channel-by-channel similarity matrices across sessions. Black horizontal lines indicate significant differences between task and residual topographies (*p <* 0.05, two-sided paired *t*-test). Gray horizontal lines indicate the noise ceiling estimated from within-task consistency. Gray triangles indicate values significantly below the noise ceiling (*p <* 0.05, one-sided *t*-test). Error bars indicate the standard error of the mean (SEM) across session pairs.

#### Topographies were adaptive across tasks, with some shared organization

Between-task consistency was substantially below the noise ceilings estimated from within-task consistency (onesided *t*-tests; mB: VWM-CDM, *t*(36) = *−*8.71, *p <* 0.001; mT: VWM-ODR, *t*(3) = *−*2.20, *p* = 0.057). However, between-task correlations were significantly greater than zero (one-sided permutation tests, *p <* 0.05), with mean correlations of *r* = 0.17 for VWM-CDM in mB and *r* = 0.20 for VWM-ODR in mT (Fig. 3c,d). Thus, although LPFC topographies were substantially reorganized across tasks, they retained a shared component of spatial organization.

This pattern indicates that task-dependent changes in LPFC topography occur on top of a partially shared spatial organization, rather than through complete reorganization of the neural landscape. Notably, this combination of task-dependent flexibility and shared organization is also consistent with our companion human fMRI studies, which identified reproducible, task-invariant organization of the human prefrontal cortex (Xiang et al., 2026, Nettekoven et al., 2026). Together, these findings suggest that prefrontal functional organization can provide a stable spatial scaffold while allowing neural population activity to be flexibly configured according to task demands.

### 2.3. Task-tuned functional topographies are more stable within tasks but more adaptive across tasks than residual activity

We next asked whether the stability and adaptability of LPFC topography differed between task-tuned responses and residual activity. Specifically, we compared topographies derived from task-tuned responses with those derived from trial-to-trial residual fluctuations in neural activity. This comparison allowed us to determine whether the temporal stability and task dependence observed above were characteristic of task-tuned neural activity or instead reflected more general properties of population activity.

#### Task-tuned topographies were more stable over time than residual topographies

Within each task, the consistency of task-tuned topographies across sessions was significantly greater than zero (one-sided permutation tests, *p <* 0.05; Fig. 4a,b). Mean consistencies were *r* = 0.43 for VWM-VWM and *r* = 0.25 for CDM-CDM in mB, and *r* = 0.64 for VWM-VWM and *r* = 0.21 for ODR-ODR in mT. More importantly, task-tuned topographies were significantly more consistent across sessions than residual topographies (two-sided paired *t*-tests; mB: VWM-VWM, *t*(54) = 3.97, *p <* 0.001; CDM-CDM, *t*(14) = 2.95, *p* = 0.01; mT: VWM-VWM, *t*(7) = 5.79, *p <* 0.001; ODR-ODR, *t*(5) = 2.60, *p* = 0.05). The dorsal array in mB showed a similar pattern (Supplementary Fig. 5). Thus, across sessions within a task, the relative organization of task-tuned responses was more stable than that of concurrent trial-to-trial fluctuations in neural activity.

**Figure 4:**
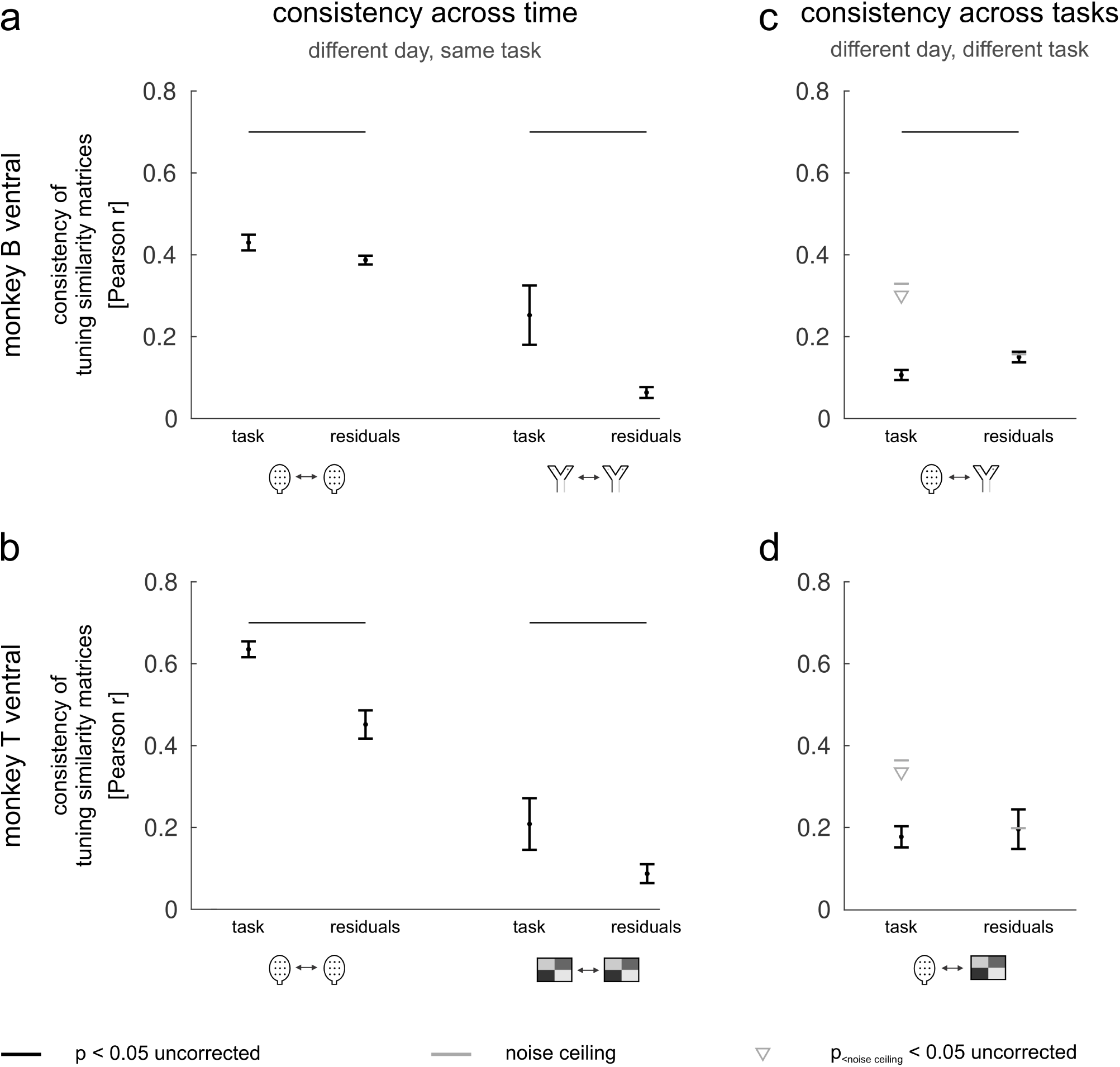
Task-tuned functional topographies are more stable within tasks but more adaptive across tasks than residual activity. **a-b)** Consistency of task-tuned functional topographies across sessions within the same task for mB (a) and mT (b). **c-d**) Consistency of task-tuned functional topographies across sessions between different tasks for mB (c) and mT (d). Task-tuned topographies are compared with topographies of residual activity to assess differences in temporal stability and task-dependent reorganization. Other conventions are as in Fig. 3.

#### Task-tuned topographies were more adaptive across tasks than residual topographies

Between tasks, task-tuned topographies remained significantly correlated (*p <* 0.05, one-sided permutation tests), with mean correlations of *r* = 0.11 for VWM-CDM in mB and *r* = 0.18 for VWM-ODR in mT. However, these correlations were substantially below the corresponding noise ceilings estimated from within-task consistency (one-sided *t*-tests; mB: VWM-CDM, *t*(36) = *−*18.04, *p <* 0.001; mT: VWM-ODR, *t*(3) = *−*7.25, *p* = 0.003; Fig. 4c,d), indicating that task-tuned topographies were not fully preserved across tasks. Moreover, between-task consistency was significantly lower for task-tuned topographies than for residual topographies in mB (VWM-CDM, *t*(36) = *−*3.03, *p* = 0.005), whereas the two measures did not differ significantly in mT (VWM-ODR, *t*(3) = *−*0.51, *p* = 0.64). The dorsal array in mB showed a similar pattern (Supplementary Fig. 5). Thus, task-tuned topographies exhibited greater task-dependent change than the topographies of concurrent trial-to-trial fluctuations.

Together, these results reveal a selective organization of task-tuned LPFC activity: task-tuned topographies were more stable across sessions within a task, yet more adaptive across tasks, than residual topographies. This suggests that the spatial organization of task-tuned neural activity is not only reproducible over time but also selectively reconfigured according to task demands.

### 2.4. Task context contributes more to topographic variation than elapsed time

The analyses above quantified topographic consistency by averaging correlations across session pairs. To complement these comparisons and quantify the relative contributions of temporal separation and task context to topographic variation, we fitted a linear regression model predicting session-pair consistency from time and task (Supplementary Results 1; Supplementary Fig. 6a). Across sessions and arrays, models including both predictors explained, on average, 87% of the variance in session-pair consistency (Supplementary Fig. 6b). Time uniquely accounted for 15% of the explainable variance, whereas task accounted for 40% (Supplementary Fig. 6c). Although the relative contributions varied across arrays, task context accounted for substantially more topographic variation than elapsed time. These results complement the session-pair analyses and further support a greater contribution of task context than temporal separation to changes in LPFC topographic organization.

### 2.5. Topographic consistency is robust to condition sampling

Finally, we tested whether our estimates of topographic consistency were robust to the specific conditions used to derive the low-dimensional projections. Using a split-half analysis with non-overlapping conditions, we asked whether tuning similarity matrices derived from disjoint subsets of conditions yielded consistent topographies. Within-task consistency remained significantly greater than zero for both same-session (Supplementary Fig. 7a) and cross-session comparisons (Supplementary Fig. 7b; one-sided *t*-tests, *p <* 0.05), demonstrating that the observed topographies were not driven by a particular subset of task conditions.

Importantly, the between-task results also reproduced the main findings. Between-task consistency remained significantly below the corresponding noise ceiling estimated from within-task consistency (Supplementary Fig. 7b; one-sided *t*-tests; mB ventral: VWM-CDM, *t*(36) = *−*15.04, *p <* 0.001; mB dorsal: VWM-CDM, *t*(36) = *−*43.68, *p <* 0.001; mT ventral: VWM-ODR, *t*(3) = *−*5.08, *p* = 0.007). These results demonstrate that the task-dependent component of LPFC topographic organization is robust to variation in condition sampling.

Together, these analyses reveal that LPFC functional topography is both stable over time and adaptive across tasks, with these properties particularly pronounced for task-tuned neural activity. Task-tuned topographies were more stable across sessions within a task than residual topographies, yet showed greater task-dependent variation across tasks. At the same time, task-tuned topographies remained significantly correlated across tasks, indicating that task-dependent reorganization occurs on top of a partially shared spatial organization. Thus, task-tuned neural activity exhibits a distinctive combination of temporal stability and taskdependent flexibility, suggesting that LPFC maintains a structured spatial organization that can be selectively reconfigured according to task demands.

### 2.6. Linking the topography of LPFC neural activity to fine-grained spatial maps

Given that feature-tuned neurons are known to cluster in populations at the spatial scale of cortical columns (Hubel and Wiesel, 1968, Merzenich and Brugge, 1973, Tanaka, 1996, Masse et al., 2017), the observed task-specific functional topographies in the LPFC (Fig. 4; Supplementary Fig. 5) may be expressed spatially by clusters of similarly tuned neurons with an adaptive task-dependent ‘fingerprint’. To quantify the degree of spatial clustering of channels with similar response preferences on the array, we computed spatial autocorrelation functions (ACFs) of channel tuning profiles (Leavitt et al., 2018, Bullock et al., 2017, Arbuckle et al., 2020, King et al., 2019). The ACFs measure the tuning similarity between channels as a function of their distance on the array. We computed ACFs for each session and task. As a measure of spatial autocorrelation, we used a cross-validated implementation of global Moran’s I that is bounded between −1 and 1, where positive values indicate similar tuning (see Methods). We found that the spatial autocorrelation between channels decreases exponentially as their distance on the array increases (Fig. 5a; Supplementary Fig. 8a,b). From the ACFs of both monkeys, we observed a positive spatial autocorrelation up to a distance of 1-2 mm across tasks (Fig. 5a).

**Figure 5:**
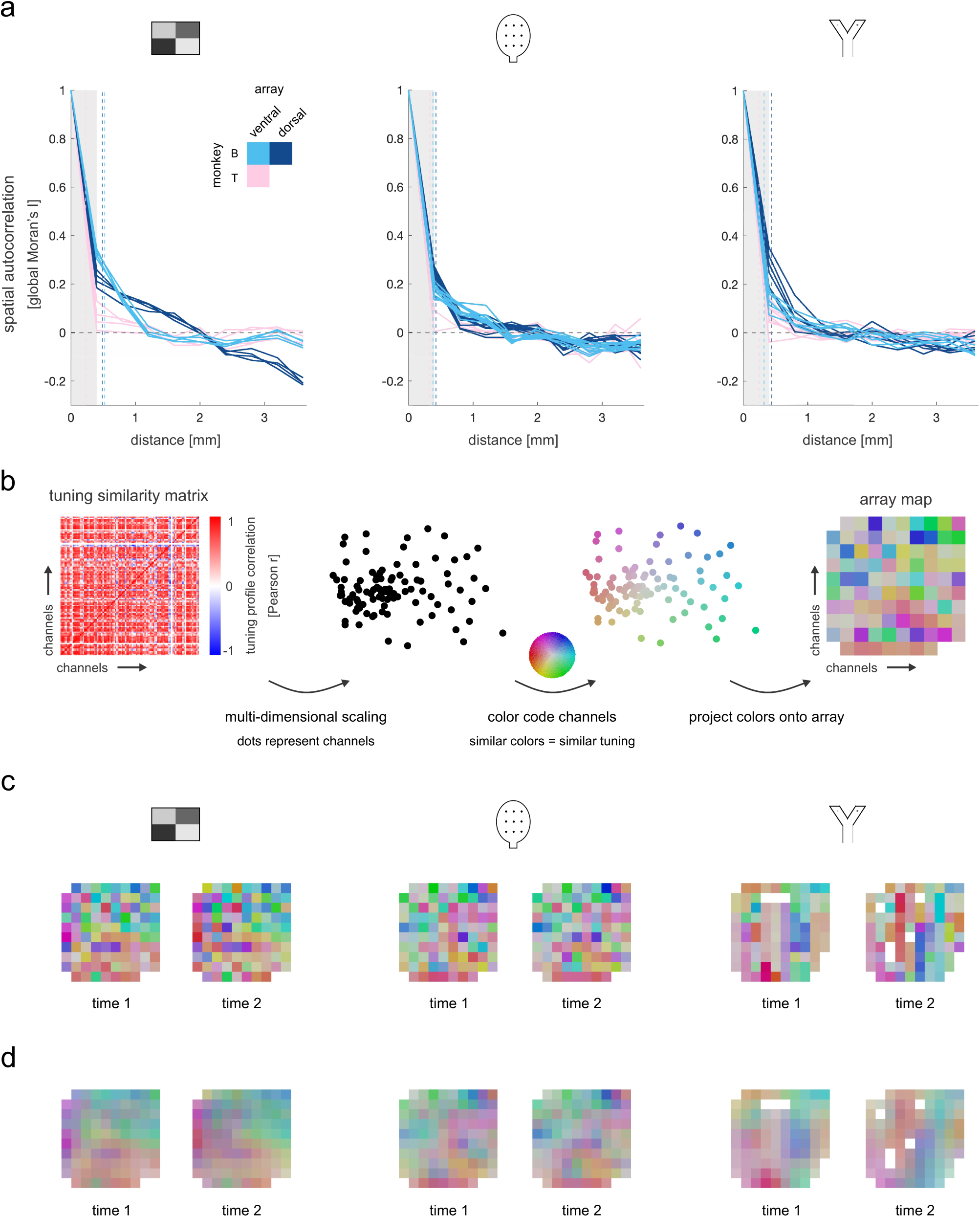
Task-tuned functional topographies in the LPFC are organized at a fine-grained spatial scale. **a)** Spatial autocorrelation functions (ACFs) for ODR, VWM and CDM tasks. Solid lines represent the ACFs for individual sessions in each task. Colors indicate arrays: light blue for the mB ventral array, dark blue for the mB dorsal array, and pink for the mT ventral array. Gray shaded areas indicate the width between two immediately neighboring channels on the array (0.4 mm). Vertical dashed lines show the median full-width-at-half-maximum (FWHM) of Laplacian functions fit to ACFs of individual sessions.1. b) Steps taken to visualize tuning similarity on the array. Channels with similar tuning profiles in a given task are similarly colored. c) Array maps for the mB dorsal array in all three tasks, using two example sessions per task. d) Array maps in panel c after smoothing with 2D Gaussian kernels whose FWHMs match that of the fitted Laplacian functions. In c and d, response profiles were aligned across sessions and tasks after dimensionality reduction using Procrustes analysis. Color mappings were defined independently for each session and indicate relative tuning-profile similarity only within that session. Similar colors across sessions or tasks do not imply similar response profiles.

These observations suggest that the task-specific functional topographies in the LPFC converge on a common spatial scale. To quantify this spatial scale, we fit Laplacian functions to the observed ACFs. Modeling the ACFs with Laplacians provides a concise description of the observed spatial decay using a limited set of parameters, allowing us to derive a scalar estimate of spatial scale. This estimate, represented by the full-width-at-half-maximum (FWHM) of the fitted Laplacians, captures the spatial extent of similarity in tuning. The FWHM is interpreted as the width of the 2-dimensional kernel needed to smooth spatially independent tuning profiles to yield the same degree of clustering we observe in the data (Diedrichsen et al., 2011). The median of the FWHMs across sessions and tasks is 369 +/-107 *µ*m (SD), suggesting the existence of fine-grained, or mesoscale, clusters of similarly tuned neural populations on the arrays.

To visualize the spatial structure of tuning similarity across the arrays, we applied two-dimensional (2D) multidimensional scaling (MDS) to the correlation distances between channel tuning profiles. Channels were color-coded according to their positions in the 2D MDS space, and these colors were projected back onto the corresponding arrays (Kiani et al., 2015) (Fig. 5b). Within each session, channels with similar tuning profiles across all conditions of a given task were assigned similar colors, allowing the spatial clustering of functionally similar channels to be visualized. Importantly, the color mapping was defined independently for each session; therefore, similar colors indicate similar response profiles only within a session and do not imply similarity across sessions or tasks. Despite this session-specific color coding, the spatial distribution of similarly tuned channels revealed clear clustering, with highly similar organization across sessions within tasks and moderate-to-weak similarity across tasks. Across all maps, these clusters exhibited a consistent spatial scale (Fig. 5c; Supplementary Fig. 8c,d). To further emphasize the spatial scale of these clusters, we smoothed the 2D MDS array maps using 2D Gaussian kernels with full-width-at-half-maximum (FWHM) values matching the empirically derived cluster sizes from the fitted Laplacian functions. Across all three tasks, this visualization highlighted the mesoscale organization of task-tuned population activity, with clusters spanning a spatial scale similar to that observed in sensory cortical areas such as V1, where mesoscale functional organization reflects tuning to stimulus features (Fang et al., 2022, Yacoub et al., 2008) (Fig. 5d).

Our findings suggest that task-tuned LPFC responses are spatially clustered at the mesoscale, similar to sensory cortical areas (Hubel and Wiesel, 1968, Merzenich and Brugge, 1973, Tanaka, 1996, Fang et al., 2022, Yacoub et al., 2008). We next asked what properties of cortical organization might shape this spatial scale. One possibility is that the afferent input projections from different regions of the brain may exhibit a spatial scale of clustering in the LPFC similar to that of the functional topographies observed in the present work. This would suggest that the structural organization of long-range inputs into the LPFC shapes the functional topographies of its task-tuned responses. To test this hypothesis, we analyzed a well-documented structural connectivity map representing the profile of afferent inputs to the macaque principal sulcus of the LPFC from the contralateral principal sulcus through callosal fibres, and from the ipsilateral parietal cortex through associational fibres (Fig. 6a,b) (Goldman-Rakic, 1984, Goldman and Nauta, 1977). The structural map overlaps the cortical patch measured with multielectrode arrays in the current study (Fig. 6a,b). To obtain an unbiased estimate of the map’s intrinsic spatial scale, we sampled 210 patches at random locations from the map, using a 4 mm *×* 4 mm patch size equivalent to the area covered by our arrays (Fig. 6b,c). For each sample, we computed a spatial ACF and corresponding FWHM using the same strategy as depicted in Fig. 5a. For both measures, we observed that the functionally derived estimates in Fig. 5a fall within the 95% confidence interval of the estimates computed across the structural map samples (Fig. 6d). These findings suggest that the structural and functional spatial scales are comparable, consistent with the idea that long-range afferent inputs may contribute to the mesoscale organization in the LPFC.

**Figure 6:**
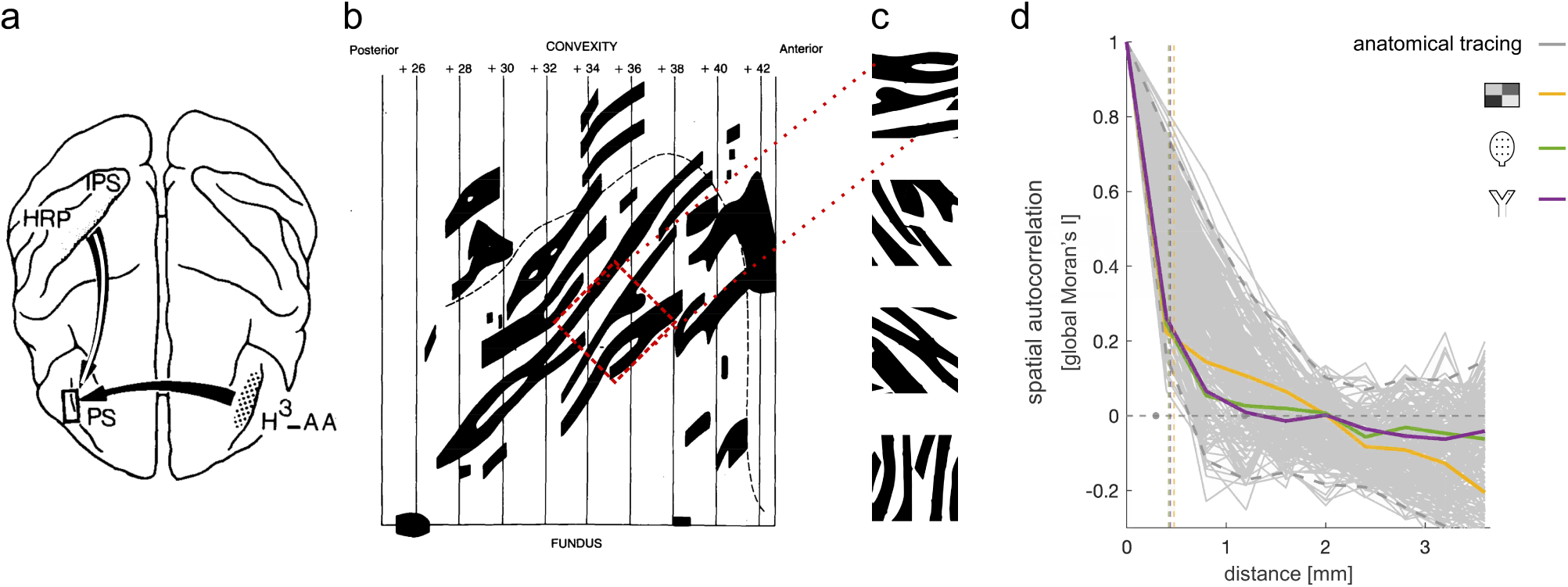
Linking task-specific functional topographies in the LPFC to fine-grained structural maps. **a)** Anatomical tracing of white matter fiber inputs to the macaque LPFC (Goldman-Rakic and Schwartz, 1982, Goldman-Rakic, 1984), adapted with permission from (Goldman-Rakic, 1984). The LPFC receives inputs through contralateral callosal fibers of the principal sulcus (PS; solid black arrow) and ipsilateral association fibers from the intraparietal sulcus (IPS; white strike arrow). *HRP* = horeradish peroxidase, *H*^3^ *− AA* = tridared amino acids. **b**) Structural map representing the spatial profile of afferent inputs to the macaque LPFC, adapted with permission from (Goldman-Rakic, 1984). The black dashed line shows the rim of the PS. Black stripes show the reconstructed terminal fields of callosal fibers in the PS, reflecting inputs from the contralateral PS. Interdigitated white stripes receive inputs from the ipsilateral IPS (Goldman-Rakic and Schwartz, 1982). The red square shows an example cortical patch sampled for our analysis. **c**) Examples of cortical patches sampled at random locations from the structural map in panel b. Each sample has a coverage of 4 mm *×* 4 mm. **d**) Functional ACFs superimposed on structural ACFs. Yellow, green and purple lines represent mean functional ACFs across sessions for the ODR, VWM and CDM tasks, respectively, for the mB dorsal array (Fig. 5a). Vertical dashed lines show the mean FWHM across sessions for each task. Solid gray lines represent structural ACFs for 210 structural map samples. Dashed gray lines show the 95% confidence interval of the structural ACFs. Gray dots placed on the zero autocorrelation line show the 95% confidence interval of the structural FWHMs.

To assess the sensitivity of these results to stripe size in the structural map, we conducted additional simulations by scaling the map to produce smaller, original, and larger stripe sizes (Supplementary Fig. 9a-c). Larger stripe sizes produced broader spatial scales in the structural ACFs, positioning the functionally derived ACFs outside the 95% confidence interval of the structural distribution at shorter distances (Supplementary Fig. 9f). Results for the smaller stripe size were less conclusive (Supplementary Fig. 9d) but did not challenge the alignment observed for the original stripe size (Supplementary Fig. 9e). These findings reinforce the link between structural and functional spatial scales by demonstrating that a broader structural scale of long-range afferent inputs is inconsistent with the functional measures.

## 3. Discussion

We used TSA to characterize the functional topography and temporal stability of task-tuned LPFC neural activity in awake, behaving macaques performing three distinct cognitive tasks (Luna et al., 2019, Roussy et al., 2021, Corrigan et al., 2022). We found that task-tuned LPFC topographies were stable across recording sessions within a task but selectively reorganized across tasks. This within-task stability was greater for task-tuned activity than for concurrent trial-to-trial residual activity, whereas task-tuned topographies showed greater task-dependent reorganization. Thus, the spatial organization of task-tuned LPFC activity was both more reproducible within a task and more sensitive to task context than that of residual activity. Despite this reorganization, task-tuned topographies retained a significant shared component across tasks. Finally, all three task-specific topographies exhibited a common spatial scale consistent with the scale of patterned afferent inputs to LPFC reported in previous anatomical tracing studies. Together, these findings suggest that LPFC functional organization comprises a partially stable spatial scaffold on which task-dependent population activity is selectively configured. Similar principles of shared and task-dependent organization were observed in our companion human fMRI studies (Xiang et al., 2026, Nettekoven et al., 2026).

TSA provides a flexible approach for characterizing the spatial organization of high-dimensional LPFC response profiles. Unlike traditional stimulus-mapping approaches (Wandell et al., 2007), TSA does not require topographic organization to be defined along a single, explicitly parameterized feature dimension. Instead, it identifies spatial patterns of similarity among channels based on their complete response profiles across task conditions (Kiani et al., 2015) (Fig. 1). This distinction is particularly relevant for LPFC, where neuronal responses can reflect mixtures of task features and their interactions (Duncan, 2001, Miller and Cohen, 2001, Rigotti et al., 2013). TSA therefore provides a means of characterizing spatial organization without prespecifying which task features should define a map, while allowing topographies derived from different tasks to be compared within a common framework. The use of a noise ceiling further provides a useful reference for interpreting the magnitude of topographic consistency (Nili et al., 2014). Nevertheless, TSA does not by itself distinguish whether spatial similarity arises from shared task tuning, intrinsic connectivity, common afferent inputs, or other sources of correlated activity. Interpreting LPFC topography therefore requires consideration of both task structure and underlying connectivity (Xiang et al., 2025).

Task mapping in LPFC presents challenges that differ fundamentally from stimulus mapping in sensory cortex. Visual stimulus dimensions such as location, orientation, and object category can be precisely parameterized, whereas task features often involve abstract relationships among rules, contexts, goals, and temporal contingencies (Duncan, 2001, Miller and Cohen, 2001, Rigotti et al., 2013, Kiani et al., 2015, Bullock et al., 2017, Leavitt et al., 2018). This distinction may help explain why the task-specific topographies observed here were not completely independent. All three tasks contained visual stimuli and shared aspects of their temporal structure, including successive cue and target phases. Some of the shared topographic organization may therefore reflect tuning to task features common across tasks, including shared temporal abstractions or task structures (Bein and Niv, 2025). Future studies could more systematically sample a high-dimensional task space in which task, stimulus, context, and behavioral demands are independently manipulated (Yang et al., 2019a, Xiang et al., 2025). Such designs would make it possible to determine whether LPFC topographies vary continuously with task similarity and to test computational accounts of prefrontal function, including biased competition (Desimone and Duncan, 1995, Schmitz and Duncan, 2018) and compositional coding (Duncan et al., 2017, Yang et al., 2019b).

The comparison between task-tuned and residual activity provides an additional constraint on the interpretation of LPFC topography. The greater within-task stability of task-tuned topographies suggests that task-related activity is embedded within a reproducible spatial organization rather than being a transient property of individual recording sessions. In contrast, the greater cross-task consistency of residual topographies indicates that some aspects of population organization are comparatively insensitive to task context. Together, these findings indicate that task context selectively reorganizes the spatial organization of task-tuned activity while leaving other aspects of population organization more conserved. Importantly, this distinction argues against a simple model in which the entire LPFC population is globally reorganized whenever task demands change (Cole et al., 2013). Instead, task context appears to modulate a pre-existing spatial organization, producing task-specific configurations of population activity.

The spatial organization observed here may reflect, in part, the heterogeneous connectivity of LPFC. Unlike sensory cortices, in which relatively local and homogeneous connectivity contributes to the formation of fine-scale stimulus maps (Yacoub et al., 2008), LPFC receives a diverse set of long-range projections as well as local intrinsic connections (Goldman-Rakic, 1988, Xu et al., 2022, Levitt et al., 1993, González-Burgos et al., 2000, Constantinidis et al., 2001, Arnsten, 2013). Anatomical tracing studies have demonstrated patterned long-range projections to LPFC from multiple cortical areas, including posterior parietal cortex and contralateral LPFC, with some projections forming elongated stripes of connectivity (Goldman-Rakic, 1988, Levitt et al., 1993). We estimated the spatial scale of these afferent patterns from a structural connectivity map reported by Goldman-Rakic and colleagues (Goldman-Rakic, 1984). The resulting scale was similar to the *∼*300–400 *µ*m spatial scale of the functional topographies observed here, despite the substantial methodological differences between the structural and functional measurements. Patterned long-range inputs, together with local horizontal connections that form patchy networks (González-Burgos et al., 2000), could therefore contribute to the spatial clustering of task-related responses. Recent multimodal studies of macaque LPFC have begun to bridge this structure-function gap using microstimulation (Xu et al., 2022) and diffusion MRI (Griffa et al., 2023). Consistent with the present findings, these studies suggest that heterogeneous long-range structural projections may provide a scaffold for functional organization on which task-dependent activity patterns are expressed (Cole et al., 2016, Xu et al., 2022).

The correspondence between the spatial scale of macaque LPFC topographies and that of patterned afferent inputs also raises the possibility that analogous functional organization could eventually be investigated in humans. The *∼*300–400 *µ*m scale observed here is challenging for conventional task-based fMRI, but submillimeter functional organization can be resolved in other cortical systems using ultra-high-field imaging (Yacoub et al., 2008). If comparable LPFC organization can be detected with high-resolution fMRI, TSA could provide a common analytical framework for linking human and macaque functional organization. Human fMRI would permit exploration of a substantially broader task space, including abstract cognitive operations that are difficult to manipulate systematically in macaques, whereas macaque electrophysiology provides the temporal and spatial resolution needed to characterize the underlying neural dynamics (Sejnowski et al., 2014). Such cross-species approaches could help establish whether the spatial scaffold identified here represents a conserved organizational principle of primate LPFC.

Several limitations should be considered. First, our ability to compare tasks was constrained by the recording schedule and by potential changes in electrode-array position across sessions. We therefore restricted comparisons to sessions separated by no more than 20 days, limiting the number of task pairs that could be examined in each animal. Second, the three tasks necessarily shared some sensory and temporal features, making it difficult to attribute cross-task similarity to a particular computational or cognitive factor. Third, the correspondence between functional topographic scale and anatomical input patterns is suggestive rather than causal. The observed spatial scale could reflect a combination of long-range afferent inputs, local connectivity, anatomical geometry, and properties of the recording array. Future experiments combining electrophysiology with direct measures of structural connectivity and systematically varied task spaces will be important for determining the mechanisms that generate and reconfigure LPFC topography (Xiang et al., 2025).

In summary, TSA revealed that task-tuned LPFC activity has a reproducible spatial organization that is stable within tasks but selectively reorganized across tasks. The greater within-task stability and greater cross-task reorganization of task-tuned activity relative to residual activity indicate that task context selectively configures, rather than globally reorganizes, LPFC population organization. At the same time, the persistence of shared topographic structure across tasks suggests that this flexibility operates within a partially conserved spatial scaffold. The common spatial scale observed across tasks, together with its correspondence to previously described patterns of long-range cortical connectivity, further suggests that anatomical organization may constrain the spatial structure within which task-dependent activity is configured. These findings provide evidence for a mode of prefrontal organization in which stable spatial structure and flexible task-dependent population coding coexist.

## 4. Methods

### 4.1. Subjects and ethics statement

We recorded LPFC neural activity in two male rhesus macaques (*Macaca mulatta*, monkey B and monkey T, 10 and 9 years old) while they were performing three different cognitive tasks across multiple measurement sessions. All training, surgery, and recording procedures conformed to the Canadian Council on Animal Care guidelines and were approved by The University of Western Ontario Animal Care Committee.

### 4.2. Behavioral tasks

#### 4.2.1. Oculomotor delayed response (ODR) task

Fig. 2a illustrates the experimental setup of the oculomotor delayed response task. Each trial began with the appearance of a fixation point at one of 16 predefined locations on a computer screen. Then a target was presented for 1000 ms before it disappeared. The monkeys were asked to maintain fixation for a variable length delay period (1400 ms −2500 ms, median = 1800 ms) and upon extinction of the fixation point, make a saccade towards the remembered target location to get a reward. More information on the ODR task can be found in (Luna et al., 2019).

#### 4.2.2. Visuospatial working memory (VWM) task

Fig. 2b illustrates the experimental setup of the visuospatial working memory task, which took place in a virtual reality environment. Within the virtual arena in the environment, targets were arranged in a 3 *×* 3 grid. The time needed to navigate between adjacent targets was *∼*0.5 s. During the cue period, a visual cue (red rectangle) was presented in one of the nine target locations for 3 seconds, then disappeared. After a 2-second delay, the monkeys navigated towards the remembered target location at a constant speed using a joystick. Upon reaching the correct target, the monkeys received a reward. More information on the VWM task can be found in (Roussy et al., 2021).

#### 4.2.3. Context-dependent decision making (CDM) task

Fig. 2c illustrates the experimental setup of the context-dependent decision making task, which was also deployed in a virtual reality environment. The task took place in a double ended Y maze, also termed “X maze” as in (Doucet et al., 2016, Gulli et al., 2020). The monkeys navigated through the X maze using a joystick. The texture of the walls, being brown “wood” or dark grey “steel”, indicated which coloured disk the monkeys should choose at the bifurcation to get a reward. In other words, the decision context was specified by the texture of the walls. We will refer to the coloured disks as goals in subsequent text. More information on the CDM task can be found in (Corrigan et al., 2022).

### 4.3. Neural recordings

Two 96-channel Utah arrays (4 mm *×* 4 mm coverage, 10 *×* 10 electrodes, spaced at 0.4 mm, 1.5 mm in length) (Blackrock Microsystems) were chronically implanted in the left LPFC in each animal. They were located anterior to the arcuate sulcus and on the ventral and dorsal side of the posterior end of the principal sulcus, targeting layers II/III of cortical areas 8A and 9/46 (see Supplementary Fig. 1 for array placement).

Neural data were recorded using a Cerebus Neural Signal Processor (Blackrock Microsystems). The neural signal was digitized (16 bit) at a sampling rate of 30 kHz. Data were online sorted to identify putative units on each recording channel in real-time. Action potential times were extracted and synchronized to task events. For all tasks, eye positions were monitored using SR Research EyeLink 1000, at a sampling rate of 500 Hz.

Both monkeys performed multiple measurement sessions for each task. Sessions were acquired on separate days. Monkey B performed 4 sessions for the ODR task, 11 sessions for the VWM task, and 6 sessions for the CDM task. Monkey T performed 4 sessions for the ODR task, 8 sessions for the VWM task, and 9 sessions for the CDM task. Three sessions were excluded from analysis in the VWM task for monkey T due to low trial numbers across experimental conditions. See Supplementary Figure 2 for further details on the included sessions. The dorsal array in monkey T was excluded from analysis due to low signal-to-noise ratio of recordings at the time of the experimental sessions, which makes it difficult to map the topography of the array.

Quality assurance data generated by the Blackrock software before each recording session indicated potential crosstalk in 4 out of the 60 sessions analyzed across monkeys and arrays. This affected only one monkey, array, and task (monkey B, dorsal, CDM) in 4 of the 6 measurement sessions for that task. To assess the robustness of the spatial scale results (Fig. 5) to potential crosstalk, we performed a conservative control analysis. For the affected sessions, we removed any shared spikes for channel pairs with more than 10% shared spikes, which impacted approximately 50 channel pairs on average per session. Results from this analysis confirmed that spike removal had minimal effects on the estimated spatial scale (Supplementary Fig. 8) and did not affect the pattern of results for the consistency analyses reported in Supplementary Figure 5.

### 4.4. Neural data analysis

#### 4.4.1. Computing task tuning profiles

To enable spatial mapping of response preferences across the array, we pooled units recorded from the same channel by summing up their spiking activities. Units recorded from the same channel exhibited similar response preferences (Supplementary Fig. 3). We refer to the pooled activity as multi-unit activity. The pooled activity reflects the activity of subpopulations of neurons within the area covered by an array channel. This area has an estimated diameter of 300 microns based on the impedance of the electrodes.

For each channel in each session in each task, we first computed trial-specific spike rates for each experimental condition and applied a square root transformation to the spike rates to account for the Poisson-like increase of variability with increasing mean firing rates (Yu et al., 2009, Arbuckle et al., 2020). We next computed trial-averaged spike rates by averaging across trials of the same experimental condition, which yielded a task tuning profile for each channel. More specifically, in the ODR task, the tuning profile for a channel was defined as the trial-averaged spike rates for the 4 quadrants (16 targets were grouped into 4 quadrants based on their location in the retinotopic reference frame, labelled 1-4 starting from bottom left, clockwise) during cue (1000 ms), delay (1400 ms −2500 ms, median = 1800 ms), and response (first 500 ms) epochs. In the VWM task, the tuning profile for a channel was defined as the trial-averaged spike rates for the 9 target locations during cue (3000 ms), delay (2000 ms), and response (first 500 ms) epochs. In the CDM task, the tuning profile for a channel was defined as the trial-averaged spike rates for the combinations of decision contexts and goal configurations in time windows before and after context onset (50 ms before, 600 ms after), goals onset (500 ms before, 300 ms after), and decision onset (500 ms before, 500 ms after). Goal configuration refers to the location of the disk colour that is associated with wood. If the colour associated with wood is on the left-hand side at the bifurcation, the trial is in configuration 1, otherwise configuration 2.

We then subtracted out the mean firing rate across all experimental conditions. The mean-centered tuning profile reflects the modulation in firing rate by the experimental conditions, providing a rich characterization of channel tuning in each task. The time windows used for computing tuning profiles were based on trial structure, monkey behavior, and neural population decoding results. Results of subsequent analyses do not critically depend on the exact time windows used.

#### 4.4.2. Computing the topography of task-tuned activity

For each array in each session in each task, we computed a channel-by-channel tuning similarity matrix. Elements in this matrix reflect the Pearson correlation of tuning profiles between channel pairs. The matrix reflects the similarity of task tuning for all channel pairs, thus capturing the similarity structure of tuning across the array. As such, it provides the basis for mapping of tuning similarity across the cortical sheet. This approach can be used to study functional topography (Kiani et al., 2015, Ito and Murray, 2023). Importantly, the tuning similarity matrices abstract from the specific experimental conditions used in a single task, which enables comparison of functional topographies between tasks.

#### 4.4.3. Computing the topography of spontaneous activity

For comparative purposes, we also analyzed trial-to-trial fluctuations about the trial averages that define the tuning profiles. Topographies based on these spontaneous fluctuations are expected to be consistent across tasks (Kiani et al., 2015, Cole et al., 2014). We partitioned the measured spiking activity in two components: task-tuned activity and spontaneous activity or residuals. For each channel in each session in each task, we computed task-tuned activity by replacing the firing rate of each trial with the mean firing rate across trials of the same experimental condition. This corresponds to a tuning profile where the trial-averaged spike rate for each condition is repeated as many times as the number of trials for that condition. We computed residuals by subtracting out the task-tuned activity from the measured spiking activity. We then computed topographies for task-tuned activity and residuals as described in the previous section, but replaced tuning profiles by task-tuned or residual activity vectors. Results reported in Fig. 4 are based on the task-tuned and residual topographies, allowing for a direct comparison between the two.

#### 4.4.4. Assessing the consistency of functional topographies across time and across tasks

To assess whether the functional topographies are consistent across time and across tasks, we computed the Pearson correlation of tuning similarity matrices across session pairs within and between tasks (see Fig. 2g). Because the matrices are symmetric, we computed correlation coefficients using the upper triangular vector of the matrix. The consistency of functional topographies within and between tasks was estimated as the mean across session pairs. To control for array shifting across time, we only included sessions spaced apart no more than 20 days for within-as well as between-task comparisons. Given that data for some tasks were acquired more than 20 days apart (see Supplementary Fig. 2), between-task comparisons are based on two out of three tasks for each monkey.

To determine whether the estimated consistencies are significantly higher than chance, we permuted channel locations on the array to simulate the null hypothesis of no consistent spatial organization. We performed 1,000 permutations, each yielding an estimate of our test statistic under the null hypothesis. If the actual consistency fell within the top 5 percent of the simulated null distribution, we rejected the null hypothesis of no consistent spatial organization. This corresponds to a one-sided test using *p <* 0.05 for significance thresholding. For between-task consistencies, we did not only test if they were higher than expected for no consistent spatial organization, we also tested if they were lower than expected for a fully consistent spatial organization. We did so by estimating a noise ceiling for between-task consistencies. For each task pair, the noise ceiling is defined as the geometric mean of the within-task consistencies, which reflects the expected maximum between-task consistency given the noise in the data (Vul et al., 2009). We compared the observed between-task consistencies against the noise ceiling using a one-sided one-sample t-test across session pairs. We compared consistencies of task-tuned and residual topographies using a two-sided paired-samples t-test across session pairs.

#### 4.4.5. Assessing the spatial scale of functional topographies

To assess the spatial scale of the functional topographies, we computed spatial autocorrelation functions (ACFs) of channel tuning profiles on the array. The spatial ACFs were computed in a cross-validated fashion to reduce the impact of spatially autocorrelated noise. Trials in a measurement session were split into two halves. Tuning profiles were estimated for each half. We used global Moran’s I as a measure of spatial autocorrelation (Moran, 1950), which is defined as:

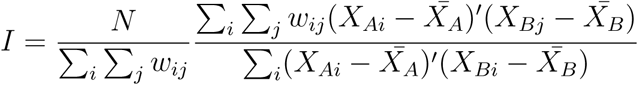

where *N* is the total number of channels on the array; *X_Ai_*is the mean-centered tuning profile for the *i^th^* channel in the first half; *X_Bj_* is the mean-centered tuning profile for the *j^th^* channel in the second half; *X*^-^ is the mean-centered tuning profile averaged across all channels, estimated for each half separately; and *w_ij_* is either 1 or 0 (if channel *i* and channel *j* are spaced within a specified spatial range, *w_ij_* equals 1; otherwise 0).

The numerator estimates the cross-validated covariances of tuning profiles between channel pairs spaced at a certain distance. The denominator estimates the cross-validated variances of tuning profiles across all the channels on the array. To account for the fact that some channels appear more frequently than others in the covariance estimates, the variance can be estimated using the following formula as in (King et al., 2019):

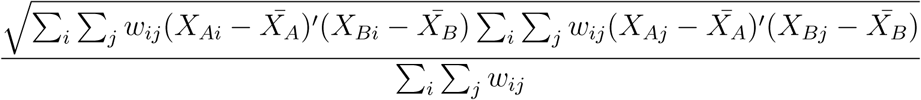

Using this formula for estimating the variance ensures that *I* is bounded between −1 and 1. A positive *I* indicates that channels spaced at the given distance have similar tuning profiles; a negative *I* indicates that channels spaced at the given distance have dissimilar tuning profiles.

To compute spatial ACFs, we systematically varied the spatial distance between channels, from including only immediately neighbouring channels (0 *<* distance *≤* 0.4 mm) to channels that are spaced apart more than one channel width but no more than two (0.4 mm *<* distance *≤* 0.8 mm), continuing these steps up and till nine channel widths (3.2 mm *<* distance *≤* 3.6 mm), and computed the spatial autocorrelation for each distance. At distance 0, the cross-validated covariances are the same as cross-validated variances, leading to a spatial autocorrelation of 1. We computed an ACF for each array in each session in each task.

To quantify the spatial scale of the functional topographies, we fit a Laplacian function to the spatial ACFs. The Laplacian function captures the exponential decay of channel tuning similarity as the distance between channels increases. The Laplacian function used is defined as follows:

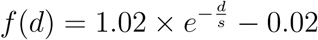

where *d ≥* 0 reflects the distances; and *s* is a fitted value, reflecting the smoothness of the curve. The full-width-at-half-maximum (FWHM) of the Laplacian curve can be computed using *s* via the following formula:

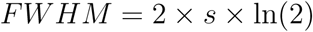

The FWHM of a fitted Laplacian function is equivalent to the FWHM of the 2-dimensional kernel required to smooth an array whose channel tuning profiles are spatially independent, to yield the degree of spatial autocorrelation we observe in the data (Diedrichsen et al., 2011). We therefore use the FWHM as an estimator of spatial scale.

#### 4.4.6. Mapping tuning similarity on the array

To visualize the spatial structure of tuning similarity on the arrays, we converted the correlations in the tuning similarity matrix to correlation distances, and applied 2D multidimensional scaling (MDS) to the distances. Channels were colour-coded based on their location in the 2D MDS space and projected back to the arrays. In the colour space, hue reflects polar angle, and saturation reflects eccentricity. Similar colours indicate similar tuning profiles. As a check, we computed the variance explained in the high-dimensional distances by the low-dimensional distances for a range of MDS dimensions (1-10). We computed the variance explained by first correlating the correlation distances between points (channels) in the original high-dimensional space with the Euclidean distances between points in the low-dimensional MDS space, and then squaring the correlation coefficients (Kiani et al., 2015). The 2D MDS space explains around 80% of the variance in the original high-dimensional space across monkeys, arrays, and tasks.

#### 4.4.7. Linking the functional topographies to structural maps

To relate the spatial scale of the observed functional topographies to prior anatomical tracing work, we repeated the spatial autocorrelation analysis on a reconstructed structural map of afferent input to macaque LPFC (Goldman-Rakic, 1984). This structural map shows the terminal field distributions of callosal fibers projecting from the principal sulcus in one hemisphere to the principal sulcus in the other hemisphere (Goldman and Nauta, 1977). The fibers terminate in a stripe-like pattern, reflecting interdigitation of the contralateral callosal fibers with associational fibers from the ipsilateral parietal cortex (Goldman-Rakic and Schwartz, 1982). The structural map suggests the existence of cortical columns in LPFC, which have been reported to have a width of 300 to 700 microns (Goldman-Rakic, 1984, Goldman and Nauta, 1977, Leichnetz et al., 1981, Goldman-Rakic and Schwartz, 1982, Bugbee and Goldman-Rakic, 1983). To assess the spatial scale of the structural map, we randomly sampled 210 cortical patches with 4 *×* 4 mm^2^ coverage from the map, simulating array placements. Sampled cortical patches were downsampled to 10 *×* 10 to match the measurement resolution of the Utah arrays used in our study. We assessed the spatial ACF for each sampled patch and plotted the distribution of ACFs across samples. We then examined whether the ACFs observed for the task-tuned topographies fall within the distribution derived from the anatomical tracing map (see Fig. 6).

## 5. Data availability

Source Data are provided with this paper. Raw data supporting the findings of this paper are available from MR, BC, RL, JMT upon reasonable requests.

## 6. Code availability

MATLAB code used for analyzing the data is available from github (https://github.com/jkderrick028/topoPFC) or from JDX.

## 7. Author contributions

JDX, TWS, JMT, MM conceptualized the work. MR designed the VWM task, collected and preprocessed the data. BC, RAG designed the CDM task, collected and preprocessed the data. RL designed the ODR task, collected and preprocessed the data. RAG performed surgical implantations. JMT contributed to experimental design. JDX analyzed the data and wrote the manuscript. MHM, LM, JD, MM provided analysis advice. JDX, TWS, JMT, MM edited the manuscript.

## 8. Acknowledgements

This work was supported by the Natural Sciences and Engineering Research Council of Canada (NSERC), the Canadian Institutes of Health Research (CIHR), Neuronex (ref. FL6GV84CKN57) grants, Ontario Graduate Scholarship, Jonathan & Joshua Memorial Graduate Scholarship in Mental Health Research, Mitacs Globalink Graduate Fellowship and McGill David G. Guthrie Fellowship. This work was further supported by BrainsCAN at the University of Western Ontario through the Canada First Research Excellence Fund (CFREF).

## 10. Supplementary information

**Supplementary Figure 1:**
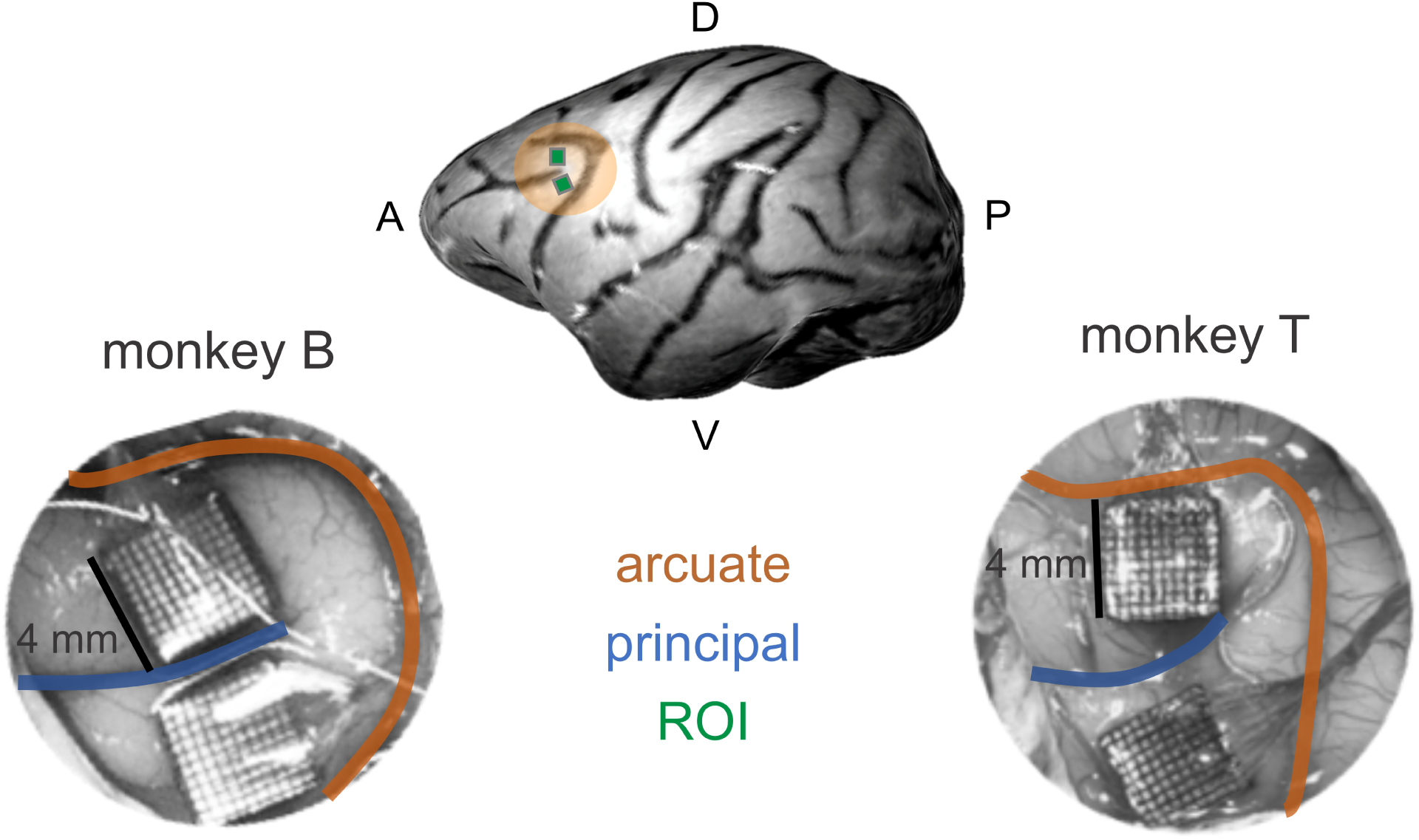
Approximate locations of the Utah arrays.

**Supplementary Figure 2:**
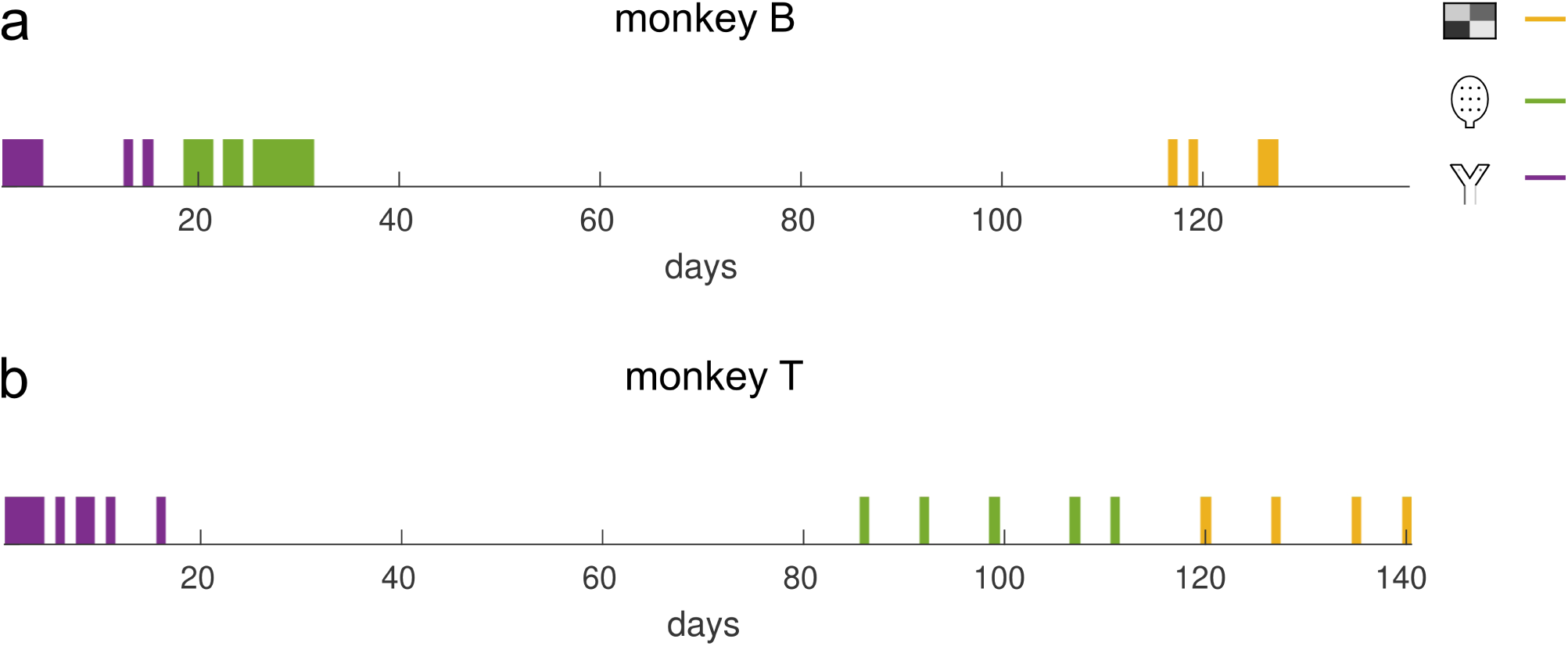
Measurement sessions for each monkey.

**Supplementary Figure 3:**
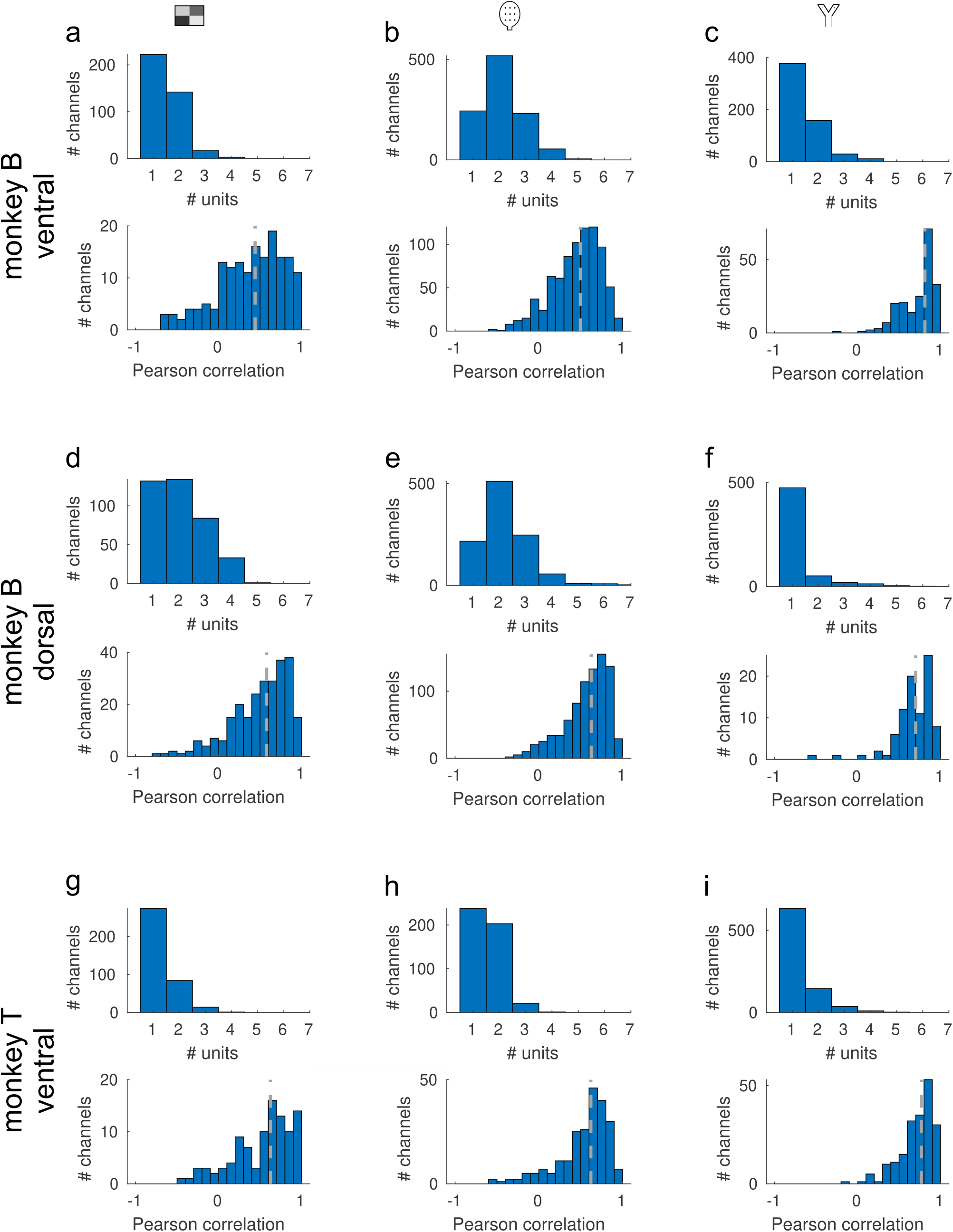
Similarity of tuning profiles of units measured by the same channel.

**Supplementary Table 1:** Number of active channels across measurement sessions for each array and task.

| array | ODR | VWM | CDM |
| --- | --- | --- | --- |
| monkey B ventral | 96.0 ± 0 (mean ± std) | 96.0 ± 0 | 95.8 ± 0.4 |
| monkey B dorsal | 96.0 ± 0 | 95.9 ± 0.3 | 93.7 ± 3.4 |
| monkey T ventral | 93.3 ± 3.2 | 92.7 ± 1.7 | 91.9 ± 0.9 |

**Supplementary Figure 4:**
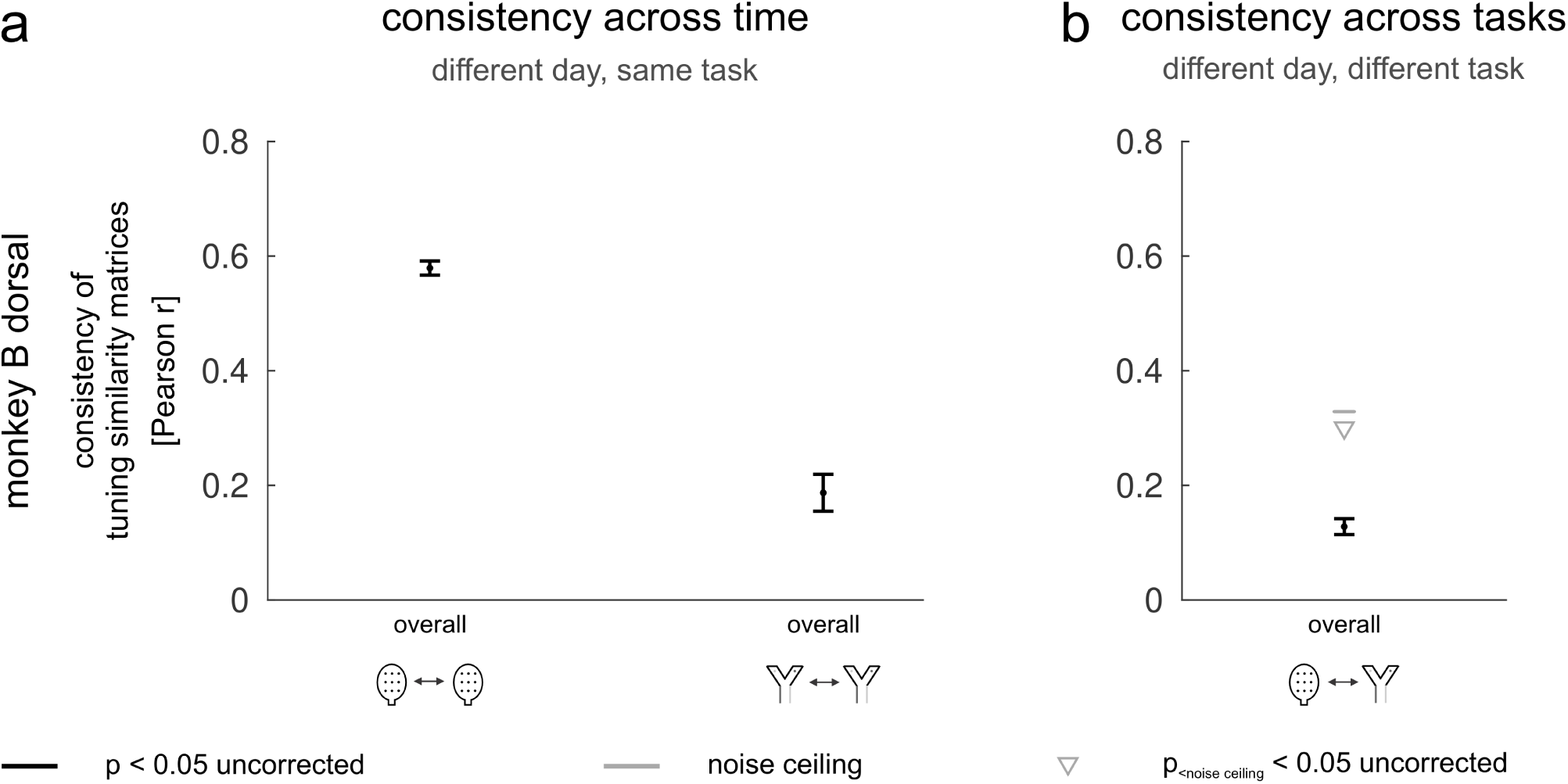
Functional topographies in the LPFC are stable across time but adaptive across tasks.

**Supplementary Figure 5:**
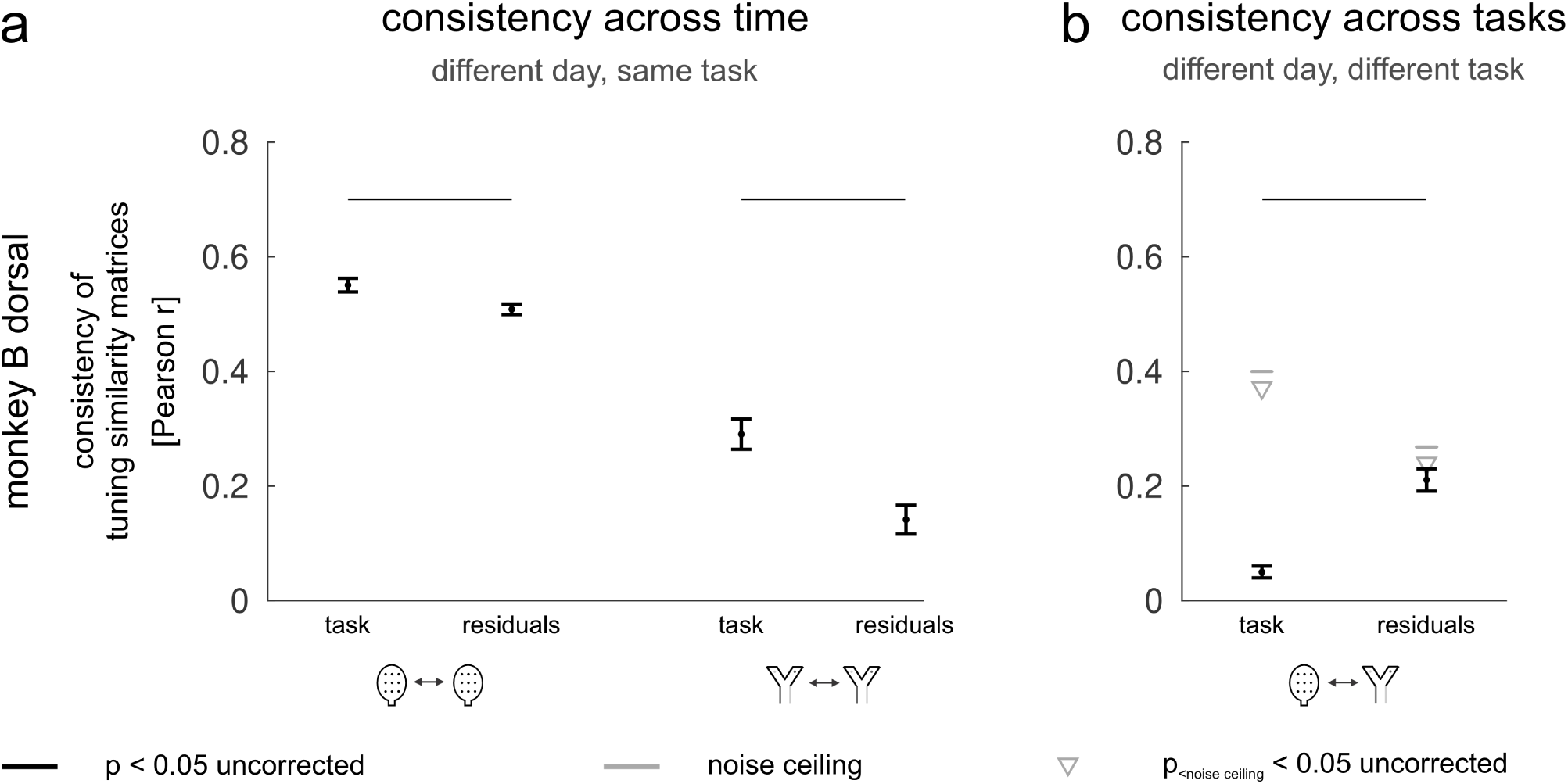
Comparing functional topographies for task-tuned responses and residuals.

### 10.1. Supplementary Results 1

To assess the relative contributions of task and time to topographic change, we quantified the unique variance explained in the consistency of tuning similarity matrices by task and by time. This was achieved using linear regression, with the full model defined as follows:

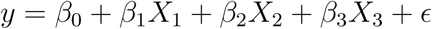

where *y* is the consistency of tuning similarity matrices between session pairs (see Supplementary Fig. 6a), *X*_1_ and *X*_2_ are indicator predictors for task (task A: *X*_1_ = 1, *X*_2_ = 0; task B: *X*_1_ = 0, *X*_2_ = 1; between task A and B: *X*_1_ = 0, *X*_2_ = 0), and *X*_3_ is a continuous predictor for time (days between sessions in a pair).

We first computed the variance explained by the full model. We next built a reduced model by removing the task predictors (*X*_1_ and *X*_2_), and computed the variance explained by the reduced model. The difference in variance explained by the full model and the reduced model is the unique variance explained by task. We used the same strategy to compute the unique variance explained by time (*X*_3_).

Results for each monkey and array are shown in Supplementary Fig. 6b,c. The full model provides a strong fit to the variance in the consistency of topography across session pairs, explaining 93% of the variance in the mB dorsal array, 78% in the mB ventral array, and 90% in the mT ventral array (Supplementary Fig. 6b). In two out of three arrays, the unique variance explained by task is substantial (Supplementary Fig. 6c): 32% of explainable variance in the mB dorsal array, 5% in the mB ventral array, and 82% in the mT ventral array. For comparison, the unique variance explained by time in these same arrays is 4%, 36%, and 5% of explainable variance, respectively. Explainable variance here refers to the variance explained by the full model. In summary, across monkeys and arrays, task uniquely accounts for 40% of the explainable variance, while time explains 15%. These results underscore the adaptability of the LPFC topography across tasks relative to its stability over time.

**Supplementary Figure 6:**
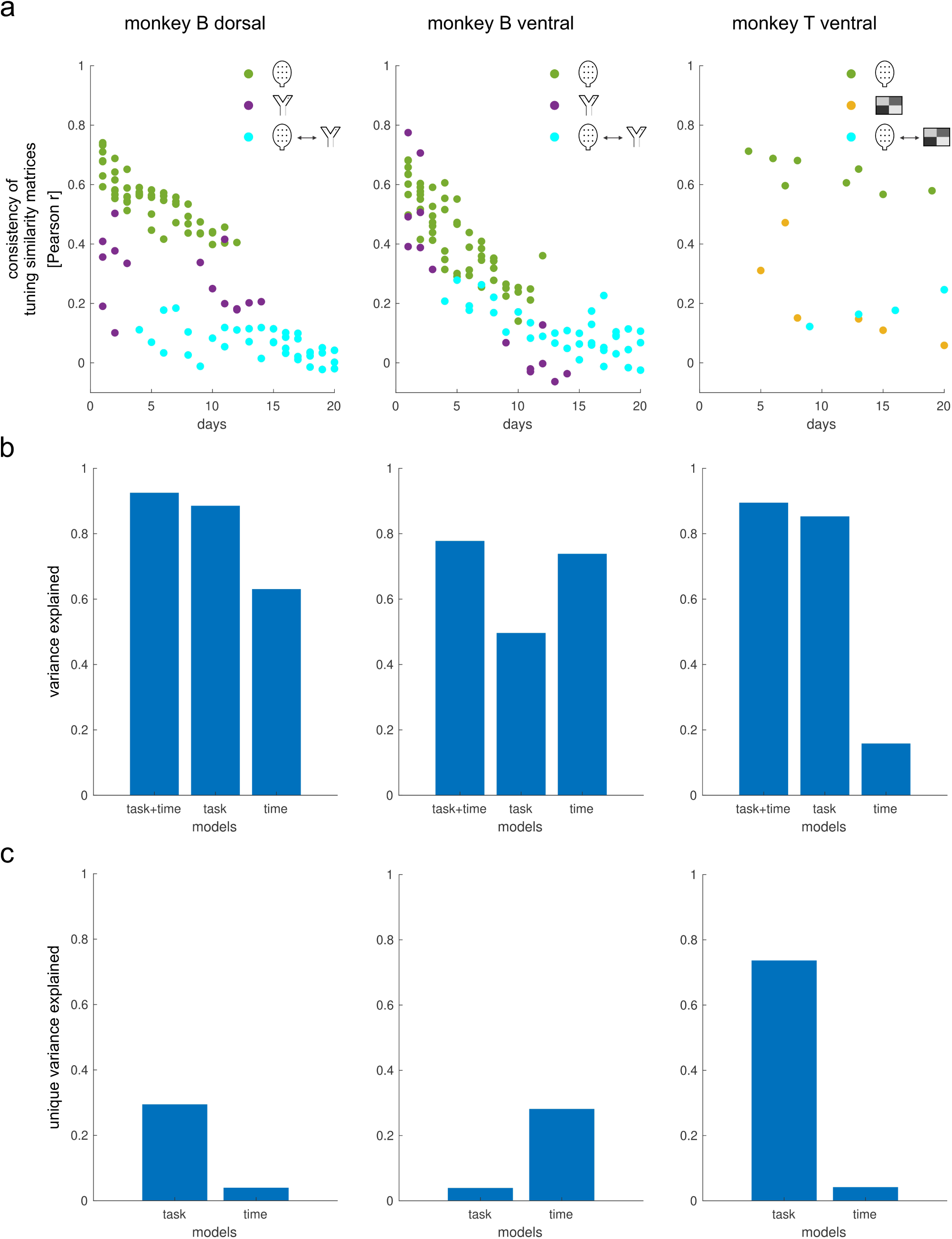
Unique contributions of task and time to topographic change in the LPFC.

**Supplementary Figure 7:**
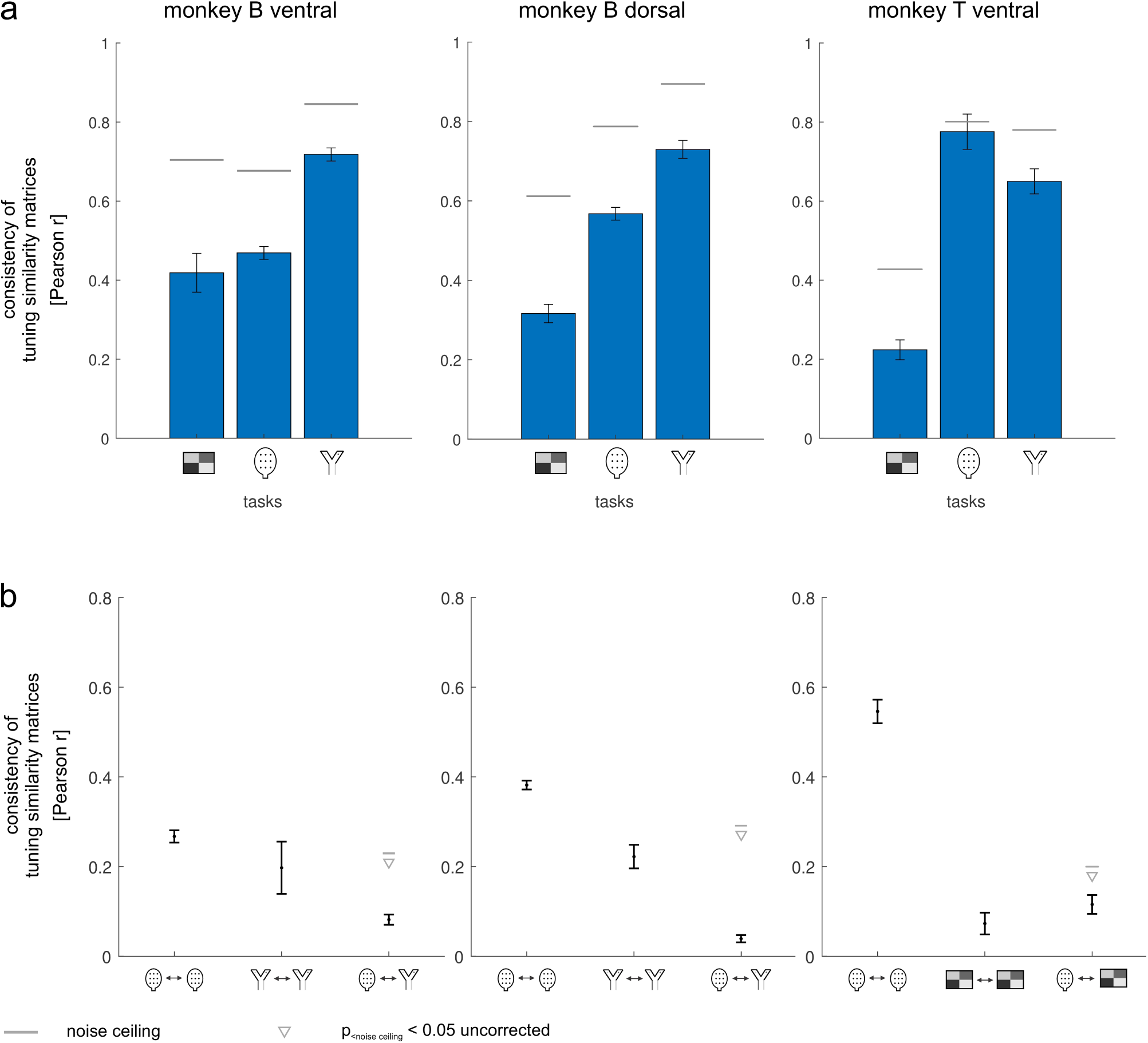
Robustness of functional topography to different low-dimensional projections of the data.

**Supplementary Figure 8:**
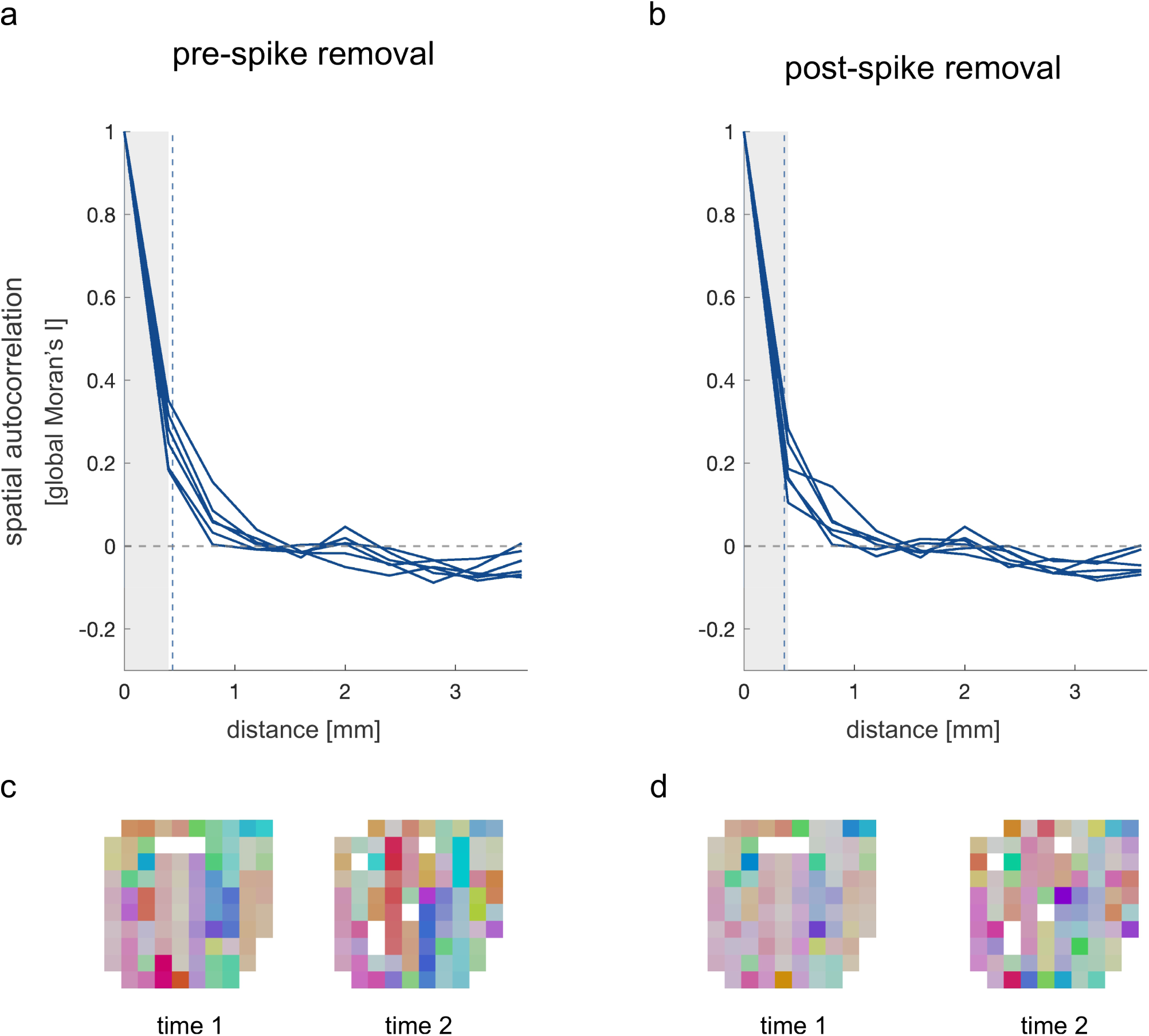
Robustness of observed spatial scale to removal of shared spikes in sessions with potential crosstalk.

**Supplementary Figure 9:**
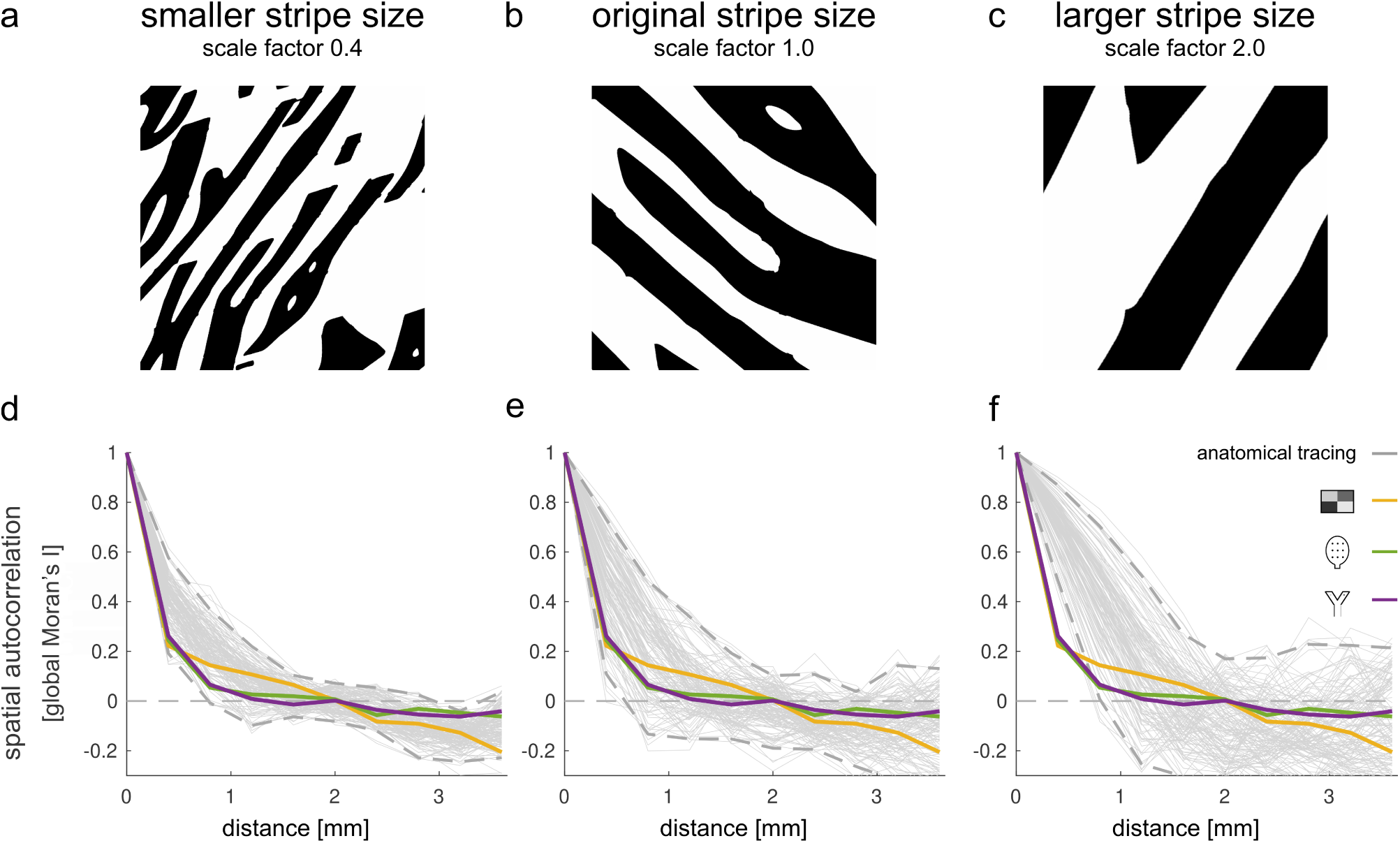
Linking task-specific functional topographies to structural maps for varying stripe sizes.

## Bibliography

S. A. Arbuckle, J. Weiler, E. A. Kirk, C. L. Rice, M. Schieber, J. A. Pruszynski, N. Ejaz, and J. Diedrichsen. Structure of Population Activity in Primary Motor Cortex for Single Finger Flexion and Extension. Journal of Neuroscience, 40(48):9210–9223, Nov. 2020. ISSN 0270-6474, 1529-2401. doi: 10.1523/JNEUROSCI.0999-20.2020. URL https://www.jneurosci.org/content/40/48/9210.

A. F. T. Arnsten. The Neurobiology of Thought: The Groundbreaking Discoveries of Patricia Goldman-Rakic 1937-2003. Cerebral Cortex, 23(10):2269–2281, Oct. 2013. ISSN 1047-3211, 1460-2199. doi: 10.1093/cercor/bht195. URL https://academic.oup.com/cercor/article-lookup/doi/10.1093/cercor/bht195.

P. Bao, L. She, M. McGill, and D. Y. Tsao. A map of object space in primate inferotemporal cortex. Nature, 583(7814):103–108, July 2020. ISSN 1476-4687. doi: 10.1038/s41586-020-2350-5. URL https://www.nature.com/articles/s41586-020-2350-5.

O. Bein and Y. Niv. Schemas, reinforcement learning and the medial prefrontal cortex. Nature Reviews Neuroscience, 26(3):141–157, Mar. 2025. ISSN 1471-0048. doi: 10.1038/s41583-024-00893-z. URL https://www.nature.com/articles/s41583-024-00893-z.

A. H. Bell, F. Hadj-Bouziane, J. B. Frihauf, R. B. H. Tootell, and L. G. Ungerleider. Object Representations in the Temporal Cortex of Monkeys and Humans as Revealed by Functional Magnetic Resonance Imaging. Journal of Neurophysiology, 101(2):688–700, Feb. 2009. ISSN 0022-3077. doi: 10.1152/jn.90657.2008. URL https://www.ncbi.nlm.nih.gov/pmc/articles/PMC2657058/.

A. A. Brewer, W. A. Press, N. K. Logothetis, and B. A. Wandell. Visual areas in macaque cortex measured using functional magnetic resonance imaging. The Journal of Neuroscience: The Official Journal of the Society for Neuroscience, 22(23):10416–10426, Dec. 2002. ISSN 1529-2401. doi: 10.1523/JNEUROSCI.22-23-10416.2002.

N. M. Bugbee and P. S. Goldman-Rakic. Columnar organization of corticocortical projections in squirrel and rhesus monkeys: Similarity of column width in species differing in cortical volume. Journal of Comparative Neurology, 220(3):355–364, 1983. ISSN 1096-9861. doi: 10.1002/cne.902200309. URL https://onlinelibrary.wiley.com/doi/abs/10.1002/cne.902200309.

K. R. Bullock, F. Pieper, A. J. Sachs, and J. C. Martinez-Trujillo. Visual and presaccadic activity in area 8Ar of the macaque monkey lateral prefrontal cortex. Journal of Neuro-physiology, 118(1):15–28, July 2017. ISSN 0022-3077. doi: 10.1152/jn.00278.2016. URL https://journals.physiology.org/doi/full/10.1152/jn.00278.2016.

M. W. Cole, J. R. Reynolds, J. D. Power, G. Repovs, A. Anticevic, and T. S. Braver. Multitask connectivity reveals flexible hubs for adaptive task control. Nature Neuroscience, 16(9): 1348–1355, Sept. 2013. ISSN 1546-1726. doi: 10.1038/nn.3470. URL https://www.nature.com/articles/nn.3470.

M. W. Cole, D. S. Bassett, J. D. Power, T. S. Braver, and S. E. Petersen. Intrinsic and Task-Evoked Network Architectures of the Human Brain. Neuron, 83(1):238–251, July 2014. ISSN 0896-6273. doi: 10.1016/j.neuron.2014.05.014. URL https://www.sciencedirect.com/science/article/pii/S0896627314004000.

M. W. Cole, T. Ito, D. S. Bassett, and D. H. Schultz. Activity flow over resting-state networks shapes cognitive task activations. Nature Neuroscience, 19(12):1718–1726, Dec. 2016. ISSN 1546-1726. doi: 10.1038/nn.4406. URL https://www.nature.com/articles/nn.4406.

C. Constantinidis, M. N. Franowicz, and P. S. Goldman-Rakic. Coding Specificity in Cortical Microcircuits: A Multiple-Electrode Analysis of Primate Prefrontal Cortex. The Journal of Neuroscience, 21(10):3646–3655, May 2001. ISSN 0270-6474, 1529-2401. doi: 10.1523/JNEUROSCI.21-10-03646.2001. URL https://www.jneurosci.org/lookup/doi/10.1523/JNEUROSCI.21-10-03646.2001.

B. W. Corrigan, R. A. Gulli, G. Doucet, M. Roussy, R. Luna, K. S. Pradeepan, A. J. Sachs, and J. C. Martinez-Trujillo. Distinct neural codes in primate hippocampus and lateral prefrontal cortex during associative learning in virtual environments. Neuron, 110(13):2155– 2169.e4, July 2022. ISSN 0896-6273. doi: 10.1016/j.neuron.2022.04.016. URL https://www.sciencedirect.com/science/article/pii/S0896627322003610.

L. Cossell, M. F. Iacaruso, D. R. Muir, R. Houlton, E. N. Sader, H. Ko, S. B. Hofer, and T. D. Mrsic-Flogel. Functional organization of excitatory synaptic strength in primary visual cortex. Nature, 518(7539):399–403, Feb. 2015. ISSN 1476-4687. doi: 10.1038/nature14182. URL https://www.nature.com/articles/nature14182.

R. Desimone and J. Duncan. Neural mechanisms of selective visual attention. Annual Review of Neuroscience, 18:193–222, 1995. ISSN 0147-006X. doi: 10.1146/annurev.ne.18.030195.001205.

J. Diedrichsen, G. R. Ridgway, K. J. Friston, and T. Wiestler. Comparing the similarity and spatial structure of neural representations: A pattern-component model. NeuroImage, 55 (4):1665–1678, Apr. 2011. ISSN 1053-8119. doi: 10.1016/j.neuroimage.2011.01.044. URL https://www.sciencedirect.com/science/article/pii/S1053811911000796.

G. Doucet, R. A. Gulli, and J. C. Martinez-Trujillo. Cross-species 3D virtual reality toolbox for visual and cognitive experiments. Journal of Neuroscience Methods, 266:84–93, June 2016. ISSN 0165-0270. doi: 10.1016/j.jneumeth.2016.03.009. URL https://www.sciencedirect.com/science/article/pii/S0165027016300152.

L. N. Driscoll, N. L. Pettit, M. Minderer, S. N. Chettih, and C. D. Harvey. Dynamic Reorganization of Neuronal Activity Patterns in Parietal Cortex. Cell, 170(5):986–999.e16, Aug. 2017. ISSN 0092-8674, 1097-4172. doi: 10.1016/j.cell.2017.07.021. URL https://www.cell.com/cell/abstract/S0092-8674(17)30828-0.

J. Duncan. An adaptive coding model of neural function in prefrontal cortex. Nature Reviews Neuroscience, 2(11):820–829, Nov. 2001. ISSN 1471-0048. doi: 10.1038/35097575. URL https://www.nature.com/articles/35097575.

J. Duncan, D. Chylinski, D. J. Mitchell, and A. Bhandari. Complexity and compositionality in fluid intelligence. Proceedings of the National Academy of Sciences, 114(20):5295–5299, May 2017. doi: 10.1073/pnas.1621147114. URL https://www.pnas.org/doi/10.1073/pnas.1621147114.

C. Fang, X. Cai, and H. D. Lu. Orientation anisotropies in macaque visual areas. Proceedings of the National Academy of Sciences, 119(15):e2113407119, Apr. 2022. doi: 10.1073/pnas.2113407119. URL https://www.pnas.org/doi/full/10.1073/pnas.2113407119.

D. J. Freedman, M. Riesenhuber, T. Poggio, and E. K. Miller. Categorical Representation of Visual Stimuli in the Primate Prefrontal Cortex. Science, 291(5502):312–316, Jan. 2001. doi: 10.1126/science.291.5502.312. URL https://www.science.org/doi/10.1126/science.291.5502.312.

W. A. Freiwald and D. Y. Tsao. Functional Compartmentalization and Viewpoint Generalization Within the Macaque Face-Processing System. Science, 330(6005):845–851, Nov. 2010. doi: 10.1126/science.1194908. URL https://www.science.org/doi/10.1126/science.1194908.

S. Fusi, E. K. Miller, and M. Rigotti. Why neurons mix: high dimensionality for higher cognition. Current Opinion in Neurobiology, 37:66–74, Apr. 2016. ISSN 0959-4388. doi: 10.1016/j.conb.2016.01.010. URL https://www.sciencedirect.com/science/article/pii/S0959438816000118.

J. Fuster. The Prefrontal Cortex. Academic Press, May 2015. ISBN 978-0-12-408060-7.

P. S. Goldman and W. J. H. Nauta. An intricately patterned prefronto-caudate projection in the rhesus monkey. Journal of Comparative Neurology, 171(3):369–385, 1977. ISSN 1096-9861. doi: 10.1002/cne.901710305. URL https://onlinelibrary.wiley.com/doi/abs/10.1002/cne.901710305.

P. S. Goldman-Rakic. Modular organization of prefrontal cortex. Trends in Neurosciences, 7(11):419–424, Nov. 1984. ISSN 0166-2236. doi: 10.1016/S0166-2236(84)80146-0. URL https://www.sciencedirect.com/science/article/pii/S0166223684801460.

P. S. Goldman-Rakic. Topography of Cognition: Parallel Distributed Networks in Primate Association Cortex. Annual Review of Neuroscience, 11(1):137–156, 1988. doi: 10.1146/annurev.ne.11.030188.001033. URL.

P. S. Goldman-Rakic and M. L. Schwartz. Interdigitation of Contralateral and Ipsilateral Columnar Projections to Frontal Association Cortex in Primates. Science, 216(4547):755– 757, May 1982. doi: 10.1126/science.6177037. URL https://www.science.org/doi/10.1126/science.6177037.

G. Gonźalez-Burgos, G. Barrionuevo, and D. A. Lewis. Horizontal Synaptic Connections in Monkey Prefrontal Cortex: An In Vitro Electrophysiological Study. Cerebral Cortex, 10(1): 82–92, Jan. 2000. ISSN 1047-3211. doi: 10.1093/cercor/10.1.82. URL.

A. Griffa, M. Mach, J. Dedelley, D. Gutierrez-Barragan, A. Gozzi, G. Allali, J. Grandjean, D. Van De Ville, and E. Amico. Evidence for increased parallel information transmission in human brain networks compared to macaques and male mice. Nature Communications, 14(1):8216, Dec. 2023. ISSN 2041-1723. doi: 10.1038/s41467-023-43971-z. URL https://www.nature.com/articles/s41467-023-43971-z.

R. A. Gulli, L. R. Duong, B. W. Corrigan, G. Doucet, S. Williams, S. Fusi, and J. C. Martinez-Trujillo. Context-dependent representations of objects and space in the primate hippocampus during virtual navigation. Nature Neuroscience, 23(1):103–112, Jan. 2020. ISSN 1546-1726. doi: 10.1038/s41593-019-0548-3. URL https://www.nature.com/articles/s41593-019-0548-3.

D. H. Hubel and T. N. Wiesel. Receptive fields and functional architecture of monkey striate cortex. The Journal of Physiology, 195(1):215–243, 1968. ISSN 1469-7793. doi: 10.1113/jphysiol.1968.sp008455. URL https://onlinelibrary.wiley.com/doi/abs/10.1113/jphysiol.1968.sp008455.

T. Ito and J. D. Murray. Multitask representations in the human cortex transform along a sensory-to-motor hierarchy. Nature Neuroscience, 26(2):306–315, Feb. 2023. ISSN 1546-1726. doi: 10.1038/s41593-022-01224-0. URL https://www.nature.com/articles/s41593-022-01224-0.

R. Kiani, H. Esteky, K. Mirpour, and K. Tanaka. Object category structure in response patterns of neuronal population in monkey inferior temporal cortex. Journal of Neurophysiology, 97 (6):4296–4309, June 2007. ISSN 0022-3077. doi: 10.1152/jn.00024.2007.

R. Kiani, C. J. Cueva, J. B. Reppas, D. Peixoto, S. I. Ryu, and W. T. Newsome. Natural Grouping of Neural Responses Reveals Spatially Segregated Clusters in Prearcuate Cortex. Neuron, 85(6):1359–1373, Mar. 2015. ISSN 0896-6273. doi: 10.1016/j.neuron.2015.02.014. URL https://www.sciencedirect.com/science/article/pii/S0896627315001282.

M. King, C. R. Hernandez-Castillo, R. A. Poldrack, R. B. Ivry, and J. Diedrichsen. Functional boundaries in the human cerebellum revealed by a multi-domain task battery. Nature Neuroscience, 22(8):1371–1378, Aug. 2019. ISSN 1546-1726. doi: 10.1038/s41593-019-0436-x. URL https://www.nature.com/articles/s41593-019-0436-x.

N. Kriegeskorte and R. A. Kievit. Representational geometry: integrating cognition, computation, and the brain. Trends in Cognitive Sciences, 17(8):401–412, Aug. 2013. ISSN 1364-6613, 1879-307X. doi: 10.1016/j.tics.2013.06.007. URL https://www.cell.com/trends/cognitive-sciences/abstract/S1364-6613(13)00127-7.

N. Kriegeskorte, M. Mur, and P. A. Bandettini. Representational similarity analysis-connecting the branches of systems neuroscience. Frontiers in Systems Neuroscience, 2, Nov. 2008a. ISSN 1662-5137. doi: 10.3389/neuro.06.004.2008. URL https://www.frontiersin.org/journals/systems-neuroscience/articles/10.3389/neuro.06.004.2008/full.

N. Kriegeskorte, M. Mur, D. A. Ruff, R. Kiani, J. Bodurka, H. Esteky, K. Tanaka, and P. A. Bandettini. Matching categorical object representations in inferior temporal cortex of man and monkey. Neuron, 60(6):1126–1141, Dec. 2008b. ISSN 1097-4199. doi: 10.1016/j.neuron.2008.10.043.

M. L. Leavitt, F. Pieper, A. J. Sachs, and J. C. Martinez-Trujillo. A Quadrantic Bias in Prefrontal Representation of Visual-Mnemonic Space. Cerebral Cortex, 28(7):2405–2421, July 2018. ISSN 1047-3211. doi: 10.1093/cercor/bhx142. URL.

G. R. Leichnetz, R. F. Spencer, S. G. P. Hardy, and J. Astruc. The prefrontal corticotectal projection in the monkey; An anterograde and retrograde horseradish peroxidase study. Neuroscience, 6(6):1023–1041, June 1981. ISSN 0306-4522. doi: 10.1016/0306-4522(81)90068-3. URL https://www.sciencedirect.com/science/article/pii/0306452281900683.

T. Lennert and J. C. Martinez-Trujillo. Prefrontal Neurons of Opposite Spatial Preference Display Distinct Target Selection Dynamics. The Journal of Neuroscience, 33(22):9520– 9529, May 2013. ISSN 0270-6474, 1529-2401. doi: 10.1523/JNEUROSCI.5156-12.2013. URL https://www.jneurosci.org/lookup/doi/10.1523/JNEUROSCI.5156-12.2013.

T. Lennert, R. Cipriani, P. Jolicoeur, D. Cheyne, and J. C. Martinez-Trujillo. Attentional Modulation of Neuromagnetic Evoked Responses in Early Human Visual Cortex and Parietal Lobe following a Rank-Order Rule. Journal of Neuroscience, 31(48):17622–17636, Nov. 2011. ISSN 0270-6474, 1529-2401. doi: 10.1523/JNEUROSCI.4781-11.2011. URL https://www.jneurosci.org/content/31/48/17622.

J. B. Levitt, D. A. Lewis, T. Yoshioka, and J. S. Lund. Topography of pyramidal neuron intrinsic connections in macaque monkey prefrontal cortex (areas 9 and 46). The Journal of Comparative Neurology, 338(3):360–376, Dec. 1993. ISSN 0021-9967. doi: 10.1002/cne.903380304.

R. Luna, M. Roussy, S. Treue, and J. C. Martinez-Trujillo. Reference Frames for Spatial Working Memory in the Lateral Prefrontal Cortex of primates. Journal of Vision, 19(10): 206, Sept. 2019. ISSN 1534-7362. doi: 10.1167/19.10.206. URL.

D. J. Margolis, H. Lütcke, K. Schulz, F. Haiss, B. Weber, S. Kügler, M. T. Hasan, and F. Helmchen. Reorganization of cortical population activity imaged throughout long-term sensory deprivation. Nature Neuroscience, 15(11):1539–1546, Nov. 2012. ISSN 1546-1726. doi: 10.1038/nn.3240.

D. A. Markowitz, C. E. Curtis, and B. Pesaran. Multiple component networks support working memory in prefrontal cortex. Proceedings of the National Academy of Sciences, 112(35): 11084–11089, Sept. 2015. doi: 10.1073/pnas.1504172112. URL https://www.pnas.org/doi/10.1073/pnas.1504172112.

N. Y. Masse, J. M. Hodnefield, and D. J. Freedman. Mnemonic Encoding and Cortical Organization in Parietal and Prefrontal Cortices. The Journal of Neuroscience, 37(25):6098–6112, June 2017. ISSN 0270-6474, 1529-2401. doi: 10.1523/JNEUROSCI.3903-16.2017. URL https://www.jneurosci.org/lookup/doi/10.1523/JNEUROSCI.3903-16.2017.

D. B. T. McMahon, I. V. Bondar, O. A. T. Afuwape, D. C. Ide, and D. A. Leopold. One month in the life of a neuron: longitudinal single-unit electrophysiology in the monkey visual system. Journal of Neurophysiology, 112(7):1748–1762, Oct. 2014. ISSN 0022-3077. doi: 10.1152/jn.00052.2014. URL https://journals.physiology.org/doi/full/10.1152/jn.00052.2014.

M. M. Merzenich and J. F. Brugge. Representation of the cochlear partition on the superior temporal plane of the macaque monkey. Brain Research, 50(2):275–296, Feb. 1973. ISSN 0006-8993. doi: 10.1016/0006-8993(73)90731-2. URL https://www.sciencedirect.com/science/article/pii/0006899373907312.

E. K. Miller and J. D. Cohen. An Integrative Theory of Prefrontal Cortex Function. Annual Review of Neuroscience, 24(1):167–202, 2001. doi: 10.1146/annurev.neuro.24.1.167. URL.

P. A. P. Moran. Notes on Continuous Stochastic Phenomena. Biometrika, 37(1/2):17–23, 1950. ISSN 0006-3444. doi: 10.2307/2332142. URL https://www.jstor.org/stable/2332142.

M. Mur, M. Meys, J. Bodurka, R. Goebel, P. A. Bandettini, and N. Kriegeskorte. Human Object-Similarity Judgments Reflect and Transcend the Primate-IT Object Representation. Frontiers in Psychology, 4, Mar. 2013. ISSN 1664-1078. doi: 10.3389/fpsyg.2013.00128. URL https://www.frontiersin.org/journals/psychology/articles/10.3389/fpsyg.2013.00128/full.

H. Muysers, H.-L. Chen, J. Hahn, S. Folschweiller, T. Sigurdsson, J.-F. Sauer, and M. Bartos. A persistent prefrontal reference frame across time and task rules. Nature Communications, 15(1):2115, Mar. 2024. ISSN 2041-1723. doi: 10.1038/s41467-024-46350-4. URL https://www.nature.com/articles/s41467-024-46350-4.

C. Nettekoven, A. Shahbazi, B. Arafat, M. Skenderija, J. D. Xiang, A. L. Pinho, and J. Diedrichsen. Multi-task fMRI outperforms resting-state fMRI for revealing task-invariant organization of the human brain, Mar. 2026. URL https://www.biorxiv.org/content/ 10.64898/2026.03.09.710558v1. ISSN: 2692-8205 Pages: 2026.03.09.710558 Section: New Results.

H. Nili, C. Wingfield, A. Walther, L. Su, W. Marslen-Wilson, and N. Kriegeskorte. A Toolbox for Representational Similarity Analysis. PLOS Computational Biology, 10(4):e1003553, Apr. 2014. ISSN 1553-7358. doi: 10.1371/journal.pcbi.1003553. URL https://journals.plos.org/ploscompbiol/article?id=10.1371/journal.pcbi.1003553.

M. Rigotti, O. Barak, M. R. Warden, X.-J. Wang, N. D. Daw, E. K. Miller, and S. Fusi. The importance of mixed selectivity in complex cognitive tasks. Nature, 497(7451):585–590, May 2013. ISSN 1476-4687. doi: 10.1038/nature12160. URL https://www.nature.com/articles/nature12160.

M. Roussy, R. Luna, L. Duong, B. Corrigan, R. A. Gulli, R. Nogueira, R. Moreno-Bote, A. J. Sachs, L. Palaniyappan, and J. C. Martinez-Trujillo. Ketamine disrupts naturalistic coding of working memory in primate lateral prefrontal cortex networks. Molecular Psychiatry, 26(11):6688–6703, Nov. 2021. ISSN 1476-5578. doi: 10.1038/s41380-021-01082-5. URL https://www.nature.com/articles/s41380-021-01082-5.

T. Sato, G. Uchida, M. D. Lescroart, J. Kitazono, M. Okada, and M. Tanifuji. Object Representation in Inferior Temporal Cortex Is Organized Hierarchically in a Mosaic-Like Structure. The Journal of Neuroscience, 33(42):16642, Oct. 2013. doi: 10.1523/JNEUROSCI.5557-12.2013. URL https://pmc.ncbi.nlm.nih.gov/articles/PMC6618530/.

T. W. Schmitz and J. Duncan. Normalization and the Cholinergic Microcircuit: A Unified Basis for Attention. Trends in Cognitive Sciences, 22(5):422–437, May 2018. ISSN 1364-6613. doi: 10.1016/j.tics.2018.02.011. URL https://www.sciencedirect.com/science/article/pii/S1364661318300470.

T. J. Sejnowski, P. S. Churchland, and J. A. Movshon. Putting big data to good use in neuroscience. Nature Neuroscience, 17(11):1440–1441, Nov. 2014. ISSN 1546-1726. doi: 10.1038/nn.3839. URL https://www.nature.com/articles/nn.3839. Number: 11.

K. Tanaka. Inferotemporal cortex and object vision. Annual Review of Neuroscience, 19: 109–139, 1996. ISSN 0147-006X. doi: 10.1146/annurev.ne.19.030196.000545.

E. Vul, C. Harris, P. Winkielman, and H. Pashler. Puzzlingly High Correlations in fMRI Studies of Emotion, Personality, and Social Cognition. Perspectives on Psychological Science, 4 (3):274–290, May 2009. ISSN 1745-6916. doi: 10.1111/j.1745-6924.2009.01125.x. URL.

B. A. Wandell, S. O. Dumoulin, and A. A. Brewer. Visual Field Maps in Human Cortex. Neuron, 56(2):366–383, Oct. 2007. ISSN 0896-6273. doi: 10.1016/j.neuron.2007.10.012. URL https://www.sciencedirect.com/science/article/pii/S089662730700774X.

J. D. Xiang, J. C. Martinez-Trujillo, T. W. Schmitz, and M. Mur. Toward task mapping of primate prefrontal cortex. Neuropsychologia, 219:109234, Dec. 2025. ISSN 0028-3932. doi: 10.1016/j.neuropsychologia.2025.109234. URL https://www.sciencedirect.com/science/article/pii/S0028393225001691.

J. D. Xiang, D. Zhi, B. Arafat, C. Nettekoven, J. Diedrichsen, and M. Mur. Individualized parcellation reveals functional boundaries in human prefrontal cortex, July 2026. URL https://www.biorxiv.org/content/10.64898/2026.07.04.736504v1. ISSN: 2692-8205 Pages: 2026.07.04.736504 Section: New Results.

R. Xu, N. P. Bichot, A. Takahashi, and R. Desimone. The cortical connectome of primate lateral prefrontal cortex. Neuron, 110(2):312–327.e7, Jan. 2022. ISSN 0896-6273. doi: 10.1016/j.neuron.2021.10.018. URL https://www.sciencedirect.com/science/article/pii/S0896627321007868.

E. Yacoub, N. Harel, and K. Ŭgurbil. High-field fMRI unveils orientation columns in humans. Proceedings of the National Academy of Sciences, 105(30):10607–10612, July 2008. doi: 10.1073/pnas.0804110105. URL https://www.pnas.org/doi/10.1073/pnas.0804110105.

G. R. Yang, M. W. Cole, and K. Rajan. How to study the neural mechanisms of multiple tasks. Current Opinion in Behavioral Sciences, 29:134–143, Oct. 2019a. ISSN 2352-1546. doi: 10.1016/j.cobeha.2019.07.001. URL https://www.sciencedirect.com/science/article/pii/S2352154619300695.

G. R. Yang, M. R. Joglekar, H. F. Song, W. T. Newsome, and X.-J. Wang. Task representations in neural networks trained to perform many cognitive tasks. Nature Neuroscience, 22(2): 297–306, Feb. 2019b. ISSN 1546-1726. doi: 10.1038/s41593-018-0310-2. URL https://www.nature.com/articles/s41593-018-0310-2.

B. M. Yu, J. P. Cunningham, G. Santhanam, S. I. Ryu, K. V. Shenoy, and M. Sahani. Gaussian-Process Factor Analysis for Low-Dimensional Single-Trial Analysis of Neural Population Activity. Journal of Neurophysiology, 102(1):614–635, July 2009. ISSN 0022-3077. doi: 10.1152/jn.90941.2008. URL https://journals.physiology.org/doi/full/10.1152/jn.90941.2008.

